# A cell-based screen identifies host-directed inhibitors and nutrient-conditional antimicrobials against intracellular *S. flexneri* dissemination

**DOI:** 10.64898/2026.11.23.768638

**Authors:** Steven Rolland, Tsuyoshi Miyake, Eric Devairakkham, Daniel A. Engel, Hervé Agaisse

## Abstract

Multidrug-resistant *S. flexneri* is a World Health Organization priority pathogen for which no universally licensed vaccine is currently available. Most antimicrobial screening platforms involve planktonic bacteria replicating in nutrient-rich medium, a condition that does not reflect the intracellular niche in which *S. flexneri* replicates and spreads and systematically excludes compounds whose activity depends on the host environment. Here, we screened the 1,581 compounds of the National Cancer Institute Diversity Set VII for inhibitors of *S. flexneri* dissemination in human colonic HT-29 monolayers by high-content microscopy. By applying sequential counterscreens, we identified six hits falling into two mechanistically distinct classes. Four compounds restricted dissemination without detectable antibacterial activity or host toxicity at the concentrations used. NSC310113 and Thiosangivamycin (NSC105827) implicate host matrix metalloprotease activity in cell-to-cell spread and host CDK2 in epithelial cell invasion. Two additional compounds, NSC45641 and NSC367428, were antibacterial only under nutrient-limited conditions and would have been described as inactive by conventional screening. NSC45641 is an adenine analog imposing a purine limitation reversible by adenine supplementation. The uracil analog NSC367428, whose activity requires the bacterial uracil phosphoribosyltransferase Upp, is consistent with perturbation of pyrimidine metabolism in both minimal medium and in HT-29 cells. Together, these findings demonstrate that cellbased dissemination screening of the NCI Diversity Set VII identified two classes of inhibitors that would likely be missed by conventional nutrient-rich medium screening, highlighting nutrient-conditional antimicrobials as promising starting points for developing treatments against intracellular pathogens.

## Introduction

The human pathogen *Shigella flexneri* is the causative agent of bacillary dysentery, or shigellosis. The disease is the second-leading cause of diarrheal mortality with 270 million cases, including more than 200,000 deaths per year worldwide ^1–3^. Young children are particularly susceptible to severe disease and repeated infections, which can contribute to growth stunting, malnutrition, impaired immune function, and increased risk of chronic inflammatory disorders later in life ^1,4^. The treatment of shigellosis is becoming increasingly challenging due to the rapid worldwide emergence of multidrug-resistant (MDR) and extensively drug-resistant (XDR) *S. flexneri* strains ^5^. Resistance to historically used first-line antibiotics, including ampicillin and trimethoprim-sulfamethoxazole, is now widespread, while resistance to newer frontline therapies such as azithromycin and ciprofloxacin continues to rise ^1^. Fluoroquinolone-resistant *S. flexneri* is classified by the World Health Organization as a priority pathogen. At the same time, no universally licensed vaccine is currently available, and the extremely low infectious dose of *S. flexneri*, estimated at only around 100 bacteria, contributes to its efficient transmission through direct contact or poor sanitation ^6^. Together, these observations highlight the urgent need to develop alternative therapeutic strategies against shigellosis.

Several non-conventional approaches have been proposed to combat MDR bacterial pathogens, including bacteriophages, antimicrobial peptides, microbiota-based therapies, and anti-virulence approaches ^7–10^. Among these, anti-virulence and host-directed therapies are particularly attractive because they impair bacterial pathogenesis without directly targeting bacterial viability, potentially reducing the selective pressure driving the emergence of antibiotic resistance. This strategy may be especially relevant for *S. flexneri*, whose pathogenesis relies extensively on the hijacking of host cellular processes to invade, survive, and disseminate within the intestinal epithelium ^11,12^. Shigellosis is characterized by bacterial invasion and dissemination throughout the colonic epithelium, leading to epithelial fenestration, massive immune cell infiltration, and bloody diarrhea ^12–16^. Efficient cell-to-cell spread is a critical determinant of tissue destruction and disease severity ^16,17^. Dissemination of *S. flexneri* depends on multiple host pathways involved in cytoskeletal remodeling, membrane trafficking, and endosomal dynamics, making these pathways attractive therapeutic targets ^18,19^. Supporting this concept, we previously demonstrated that pharmacological inhibition of the host lipid kinase PIK3C3/hVPS34, which promotes a key step of cellto-cell spread, significantly reduces *S. flexneri* dissemination in epithelial cells and within the colonic mucosa of the infant rabbit model ^17^. In addition, several *S. flexneri* mutants impaired in dissemination exhibit markedly attenuated virulence *in vivo* ^16,20,21^. Together, these studies highlight the therapeutic potential of targeting host-dependent mechanisms required for *S. flexneri* dissemination.

An underappreciated challenge in antimicrobial discovery against *S. flexneri* and other intracellular bacteria is that conventional screening approaches fail to account for the intracellular lifestyle and the environment in which the pathogen replicates. *S. flexneri* is a facultative intracellular bacterium that invades non-phagocytic cells, such as colonic epithelial cells, escapes the entry vacuole, replicates in the cytosol, and spreads into adjacent cells, resulting in tissue colonization ^14,16^. Conventional screening assays are usually based on planktonic bacteria replicating in rich laboratory media, a condition that differs from the intracellular niche for many pathogens including *S. flexneri*. Limitations stem from at least two main factors. First, compounds that display no activity in rich media, but exhibit antimicrobial activity in certain conditions are excluded from these screens. Secondly, compounds that do not penetrate host cells do not engage their intracellular bacterial target, thereby limiting their efficacy against intracellular pathogens. This does not reflect what is observed in rich medium where the contact is direct and bacterial replication conditions are optimal, leading to a misleading evaluation of the antibacterial activity. Cell-based phenotypic assays reflecting the intracellular lifestyle become an interesting approach to identifying new types of molecules. Cell-based screening has recently been applied to *S. flexneri* on a considerable scale. A screen of 6,454 compounds identified 31 inhibitors of *S. flexneri* replication in enterocyte-like cells and showed that antibacterial activity depends strongly on the composition of the medium ^22^. High-throughput screening campaigns assessing bacterial survival in macrophages and intracellular replication in three-dimensional Caco-2 cells have screened 1.65 million and 518,282 compounds, respectively ^23,24^.

Here, we designed a novel screening strategy to identify compounds targeting post-entry stages of infection, with a particular focus on bacterial spread from cell to cell. We used the approach to conduct a microscopy-based phenotypic screen of the 1,581 compounds comprising the National Cancer Institute (NCI) Diversity Set VII to identify inhibitors of intracellular dissemination of *S. flexneri* in human colonic epithelial HT-29 cells. A series of sequential counter-screening assays, assessing bacterial growth in nutrient-rich and nutrient-limited media as well as host cell viability, allowed us to exclude compounds displaying direct antibacterial activity or cytotoxic effects on host cells. We provide initial characterization of the activity of prioritized hits.

## Results

### High-content screening of the NCI diversity set VII identifies inhibitors of *S. flexneri* cell-to-cell spread in an HT-29 cell monolayer

*S. flexneri* is a facultative intracellular bacterium characterized by its ability to enter into nonphagocytic cells, such as epithelial cells, and spread into adjacent cells. We performed a high-content screening using the NCI Diversity Set VII consisting of 1,581 compounds designed and selected from a pool of more than 140,000 compounds. The compounds were added at 20 µM (final concentration) onto a 4-day HT-29 cell monolayer for one-hour prior to infection. CFP-expressing *S. flexneri* were incubated with cells for one hour before adding gentamicin to kill extracellular bacteria and IPTG to induce CFP expression. At 8 hpi, the infection foci generated by CFP-expressing *S. flexneri* were analyzed by fluorescence microscopy (Fig 1A; Supplementary Fig 1). We identified 61 primary hits showing a defect in infection foci size (Fig 1B, Supplementary table 1). After visual inspection of the images, compounds were excluded based on (i) autofluorescence or precipitation of the compounds, (ii) absence of infection foci, and (iii) obvious compound-mediated cell toxicity challenging the cell monolayer integrity. This visual quality control excluded 27 compounds (Fig 1C, Supplementary. Figure 2, Supplementary. Table 1), and independent samples of the remaining 34 compounds were re-ordered (Supplementary Table 2).

**Figure 1:**
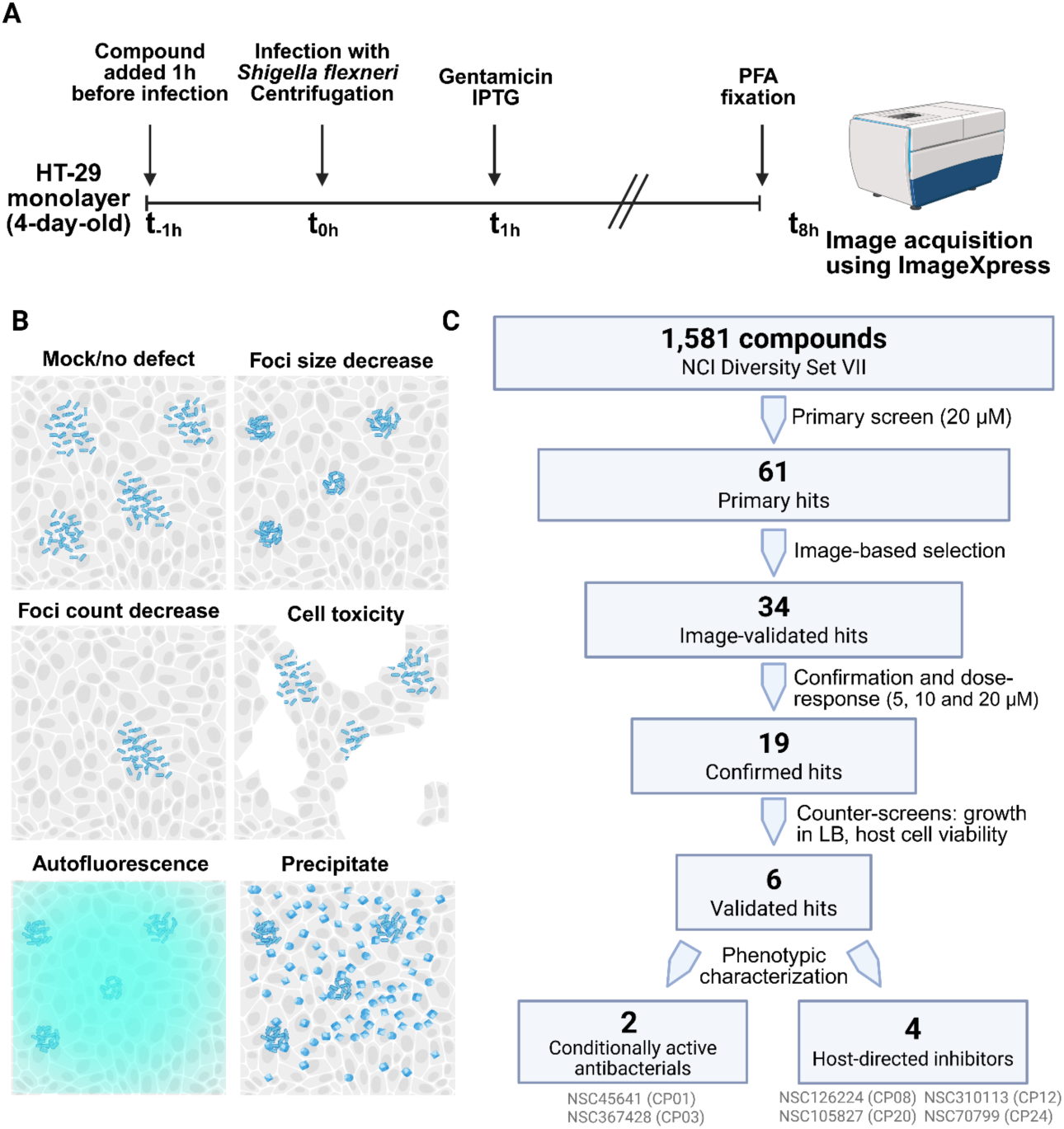
Workflow of the selection of hits, focusing on Host-directed therapeutics to inhibit intracellular *Shigella flexneri*. (A) Representation of the infection process of CFP-expressing Shigella in an HT-29 epithelial cell monolayer. (B) Cartoon representing the possible phenotypic outcomes from our approach. (C) Flow of the different steps leading to hit selection.

The 34 compounds were re-tested in dose response 5 µM, 10 µM and 20 µM; Supplementary. Table 2), and the same quality control criteria were re-applied to the resulting images. On the basis of reproducibility of the original phenotype, autofluorescence and cytotoxicity, we selected 19 compounds for further characterization (Table 1).

**Table 1:** Cytotoxicity and anti-*Shigella* activity of selected compounds identified in the primary screen prior to counterselection. The table summarizes the nineteen compounds identified in the initial screen before counterselection. For each compound, the studyspecific identifier (CP), official NSC code, and generic name, when available, are provided. Cell viability percentages following treatment with 31.6 µM of each compound for 8 h and 24 h are reported, along with the estimated IC₅₀ values at both time points, based on the viability assay results presented in Supplementary Table 3. TUNEL assay results and effects on bacterial growth are also indicated for the six compounds selected for further characterization (highlighted in white).

|  | NSC# | Generic name | 8h (screen time point) |  |  | 24h |  |  | TUNEL Assay (% nuclei at 24h) | Anti-Shi-gella (Fig 2) |
|  |  |  | IC50 (µM) | %viability at 31.6 µM | Toxic at 31.6 µM | IC50 (µM) | %viability at 31.6 µM | Toxic at 31.6 µM |  |  |
|---|---|---|---|---|---|---|---|---|---|---|
| **CP01** | 45641 | MYRA-B | >100 | 126.2 ±6.1 | nontoxic | >100 | 104.2±5.7 | nontoxic | 2.05±0.19 n.s | Cond. |
| **CP02** | 303603 | Nifuratrone | >100 | 103.0±5.0 | nontoxic | >100 | 87.1±1.8 | nontoxic | N/A | **Yes** |
| **CP03** | 367428 |  | >100 | 99.1±3.7 | nontoxic | >100 | 106.2±4.4 | nontoxic | 1.82±0.57 n.s | Cond. |
| **CP05** | 59984 |  | >100 | 91.4±3.3 | nontoxic | >100 | 87.6±0.5 | nontoxic | N/A | **Yes** |
| CP08 | 126224 |  | >100 | 101.4±1.4 | nontoxic | **57.3** | 85.6±3.5 | nontoxic | 2.94±1.80 n.s | No |
| **CP09** | 207895 | XI-006 | >100 | 108.7±2.7 | nontoxic | **18.1** | 14.4±1.8 | toxic | N/A | **Yes** |
| CP12 | 310113 |  | >100 | 101.3±1.1 | nontoxic | >100 | 98.6±1.8 | nontoxic | 1.96±0.18 n.s | No |
| CP14 | 33353 |  | >100 | 92.8±2.2 | nontoxic | **23** | 29.7±1.4 | toxic | N/A | No |
| **CP16** | 30205 |  | >100 | 95.4±1.4 | nontoxic | **70.6** | 88.9±0.6 | nontoxic | N/A | **Yes** |
| **CP17** | 59814 | Oxyphenisatin | >100 | 70.6±6.9 | toxic | **N.A** | 47.3±1.5 | toxic | N/A | No |
| CP20 | 105827 | FLIX1; Thiosangi-vamycin | >100 | 99.2±0.9 | nontoxic | >100 | 83.8±0.8 | nontoxic | 5.42±0.19 *** | No |
| **CP21** | 133114 | 5’-Sulfamoyladeno-sine | >100 | 96.1±3.7 | nontoxic | **<1** | 13.6±2.9 | toxic | N/A | **Yes** |
| **CP22** | 339316 |  | >100 | 97.6±0.8 | nontoxic | **68.1** | 60.0±1.1 | toxic | N/A | Yes |
| CP24 | 70799 |  | >100 | 99.3±0.6 | nontoxic | >100 | 96.8±1.2 | nontoxic | 1.69±0.19 n.s | No |
| **CP26** | 309401 |  | **91.9** | 104.5±1.1 | nontoxic | **43.1** | 74.1±1.4 | toxic | N/A | **Yes** |
| **CP29** | 325014 | Bactobolin | >100 | 109.0±0.5 | nontoxic | >100 | 93.7±0.4 | nontoxic | N/A | **Yes** |
| **CP30** | 65238 |  | **32.7** | 53.3±5.1 | toxic | **16.8** | 5.9±1.9 | toxic | N/A | **Yes** |
| **CP32** | 277184 |  | >100 | 110.6±1.4 | nontoxic | **53.4** | 72.0±0.4 | toxic | N/A | **Yes** |
| **CP34** | 19990 | Streptovaricin C | >100 | 100.0±0.1 | nontoxic | >100 | 95.4±3.1 | nontoxic | N/A | **Yes** |

### Selection of compounds that do not affect bacterial growth

Compounds that directly impair bacterial replication would be expected to reduce infection foci size independently of any effect on cell-to-cell spread. To identify those compounds that may display antibiotic activities, we cultured *S. flexneri* in the presence of 20 µM of each compound in liquid LB broth at 37°C. Bacterial growth was measured by optical density (OD600nm) during the mid-exponential phase of growth. Eleven compounds showed growth-inhibitory effects, including five that partially affected growth in LB medium (Fig 2A). Interestingly, CP03 (NSC367428) failed to inhibit growth in rich media, despite structural similarities with the known antimetabolite 5-fluorouracil (5-FU). Since 5-FU interferes with nucleotide metabolism, an effect that can be overlooked in rich media, such as LB broth, we re-evaluated growth in a defined minimal medium for *S. flexneri* ^25^. Under these nutrient-restricted conditions, CP01 (MYRA-B/NSC45641) and CP03 significantly inhibited *S. flexneri* growth (Fig 2B). CP01, an adenine-related structure previously shown to induce Myc-dependent apoptosis, likely interferes with bacterial purine metabolism ^26^.

**Figure 2:**
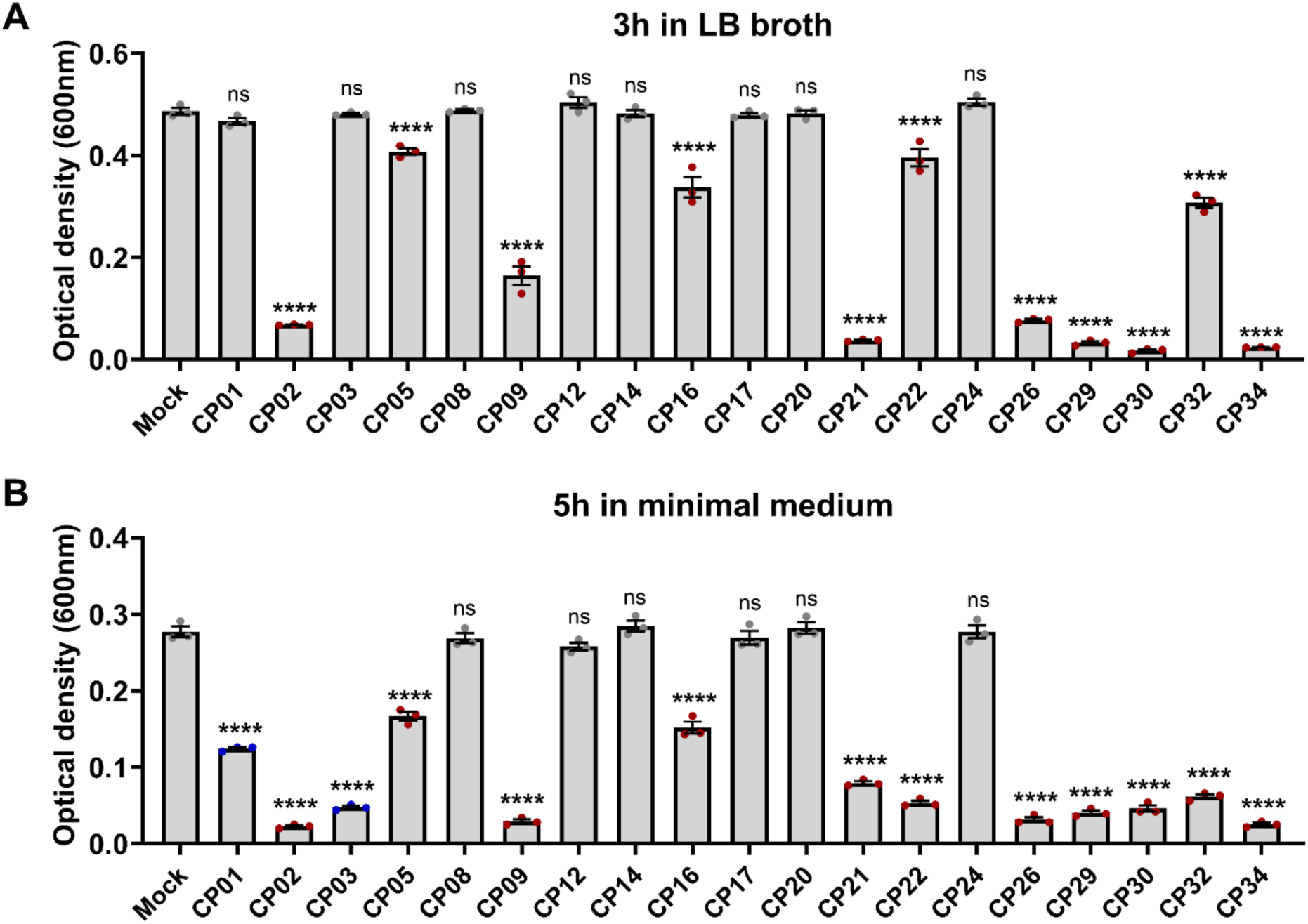
Effect of the nineteen selected compounds on *Shigella flexneri* growth. Bacterial growth was assessed by measuring optical density at 600 nm (OD₆₀₀) after 3 h of incubation in LB medium (A) or 5 h in minimal medium (B), in the presence of nineteen selected compounds from the NCI Diversity Set VII (CP01–CP34) at a final concentration of 20 µM. Each dot represents an independent biological replicate. Error bars indicate the standard error of the mean (SEM). Statistical significance was determined by one-way ANOVA followed by Dunnett’s multiple-comparisons test. ns, not significant; ****, p < 0.0001.

### Assessment of compound cytotoxicity

To confirm that the observed phenotypes were not due to negative effects on the host cell physiology, we assayed cell viability by quantifying ATP production across a semi-logarithmic range (1.0–100 µM) at both 8 and 24 hours. According to ISO 10993-5, a reduction in cell viability to below 70% of the untreated control is considered as indicative of cytotoxicity ^27^. We applied a more stringent threshold of 80% to define the non-cytotoxic range, excluding compounds that caused even moderate reductions in cell viability. This margin distinguishes minor metabolic fluctuations from biologically relevant cellular distress. After 8 hours of exposure, corresponding to the infection time window of our screen, 17 of the 19 compounds maintained a mean viability over 80 % at 31.6 µM, a concentration exceeding the 20 µM used in the primary screen. Only CP17 (NSC59814 or oxyphenisatin) and CP30 (NSC65238) affected cell viability (Table 1). Since cell viability remained unaffected at 31.6 µM at both time points for 11 of these compounds, we reasoned that the reduction in *S. flexneri* infection focus size observed at the lower concentration of 20 µM was unlikely to result from indirect effects of host cell death. To evaluate potential long-term effects relevant to therapeutic applications, viability was also assessed following a 24-hour compound exposure. Under these conditions, eight compounds, including CP17 and CP30, affected cell viability and were excluded from further analysis (Table 1; Supplementary Table 3). Of the 19 compounds, six did not cause a *S. flexneri* growth defect in rich medium nor affected HT-29 cell viability. To verify that these six compounds were not triggering cell death, we performed an additional validation using a TUNEL assay highlighting DNA breakage on an HT-29 cell monolayer treated for 24 hours with 20 µM of each of these compounds. Five compounds gave a proportion of TUNEL-positive nuclei comparable to the mock-treated control (Table 1, Supplementary Table 3). CP20 caused a small but statistically significant increase. Because this increase remained modest and did not reflect massive cell death, CP20 was not excluded and was retained for further characterization.

In conclusion, six compounds were prioritized for further study including four that presented no direct antimicrobial pressure in rich or nutrient-limited medium, suggesting a robust host-targeted mode of action (CP08, CP12, CP20 and CP24; Table 1), and two compounds presenting a growth defect under nutrient-depleted conditions(CP01 and CP03; Table 1).

### CP08 (NSC126224) restricts *S. flexneri* dissemination independently of canonical TLR signaling

We identified NSC126224 (CP08) as one of four compounds that significantly reduced infection foci size in HT-29 monolayers (Supplementary tables 1 and 2). Treatment with NSC126224 led to a marked dose-dependent decrease in *S. flexneri* infection foci size without affecting bacterial invasion (Supplementary Figs 3A and 3G). As previously described, NSC126224 did not significantly affect ATP levels after 8 h of treatment, corresponding to the duration of the infection assay (Table 1; Supplementary table 3). A reduction in ATP levels was observed after 24 h at a higher dose (IC50 = 57.3 µM), suggesting a delayed impact on cellular metabolism. In addition, NSC126224 did not present direct antibacterial activity against *S. flexneri* (Fig. 2A, Fig 2B). A previous study reported NSC126224 as an antagonist of TLR4 and TLR2 signaling pathways through suppression in LPS-induced production of nuclear factor kappaB (NF-kB), tumor necrosis factor (TNFa), Interleukin-1beta, and nitric oxide (NO) ^28^. To test whether inhibition of Toll-like receptor signaling could be responsible for the observed decrease in size of the infection foci formed in HT-29 cells, we incubated cells for 1 hour before infection with the TLR4 inhibitor TAK-242 (Supplementary Figs 3B, 3H), the TLR2 inhibitor C29 (Supplementary Figs 3C, 3I), the NFκB pathway inhibitor BAY 11-7082 (Supplementary Figs 3D, 3J), inducible nitric oxide synthase (iNOS) inhibitor 1400W (Supplementary Figs 3E, 3K), or the nitric oxide synthase (NOS) inhibitor L-NAME (Supplementary Figs 3F, 3L) ^29–32^. None of these compounds recapitulated the reduction in infection foci size observed with NSC126224. Together, these results suggest that NSC126224 restricts *S. flexneri* cell-to-cell spread through a mechanism that is independent of canonical TLR signaling. This suggests that NSC126224 targets a distinct host cellular process required for efficient bacterial dissemination.

### Metalloprotease activity contributes to *S. flexneri* cell-to-cell spread

We identified NSC310113 (CP12) as one of four compounds that significantly reduced infection foci size in HT-29 monolayers (Supplementary tables 1 and 2). Treatment with NSC310113 led to a dose-dependent decrease in the size of *S. flexneri* infection foci without affecting bacterial invasion (Fig 3A, 3B). As previously described, this compound did not significantly affect ATP levels either following 8 h of treatment, corresponding to the duration of the infection assay, or after a longer incubation time (Table 1; Supplementary table 3). In addition, it did not exhibit direct antibacterial activity against *S. flexneri* (Fig. 2A, Fig 2B). We observed a dosedependent decrease of *S. flexneri* infection foci size (Fig 3A) without affecting the infection foci count (Fig 3B). To our knowledge, CP12 does not have any characterized activity described in the literature. However, CP12 structure (ethyl 4oxo-3,4,5,6,7,8-hexahydro[1]benzothieno[2,3-d]pyrimidine-2-carboxylate) is identical to a compound disclosed in WO2008063667, a patent covering thieno-fused pyrimidin-4-ones as inhibitors of MMP-13, MMP-8, MMP-3 and MMP-2 metalloproteases ^33^. We therefore tested whether the broad-spectrum metalloprotease inhibitor Ilomastat (GM6001) would reproduce the same phenotype ^34^. Ilomastat decreases the foci size in a dose-dependent manner without affecting the foci count (Figs 3C, 3D). To exclude a direct antibacterial effect, Ilomastat was tested in bacterial growth assays. No growth defects were observed compared to control conditions, confirming that the observed phenotypes are host-mediated (Fig 3E). These results suggest a role of matrix metalloproteinases in *S. flexneri* dissemination into adjacent cells.

**Figure 3:**
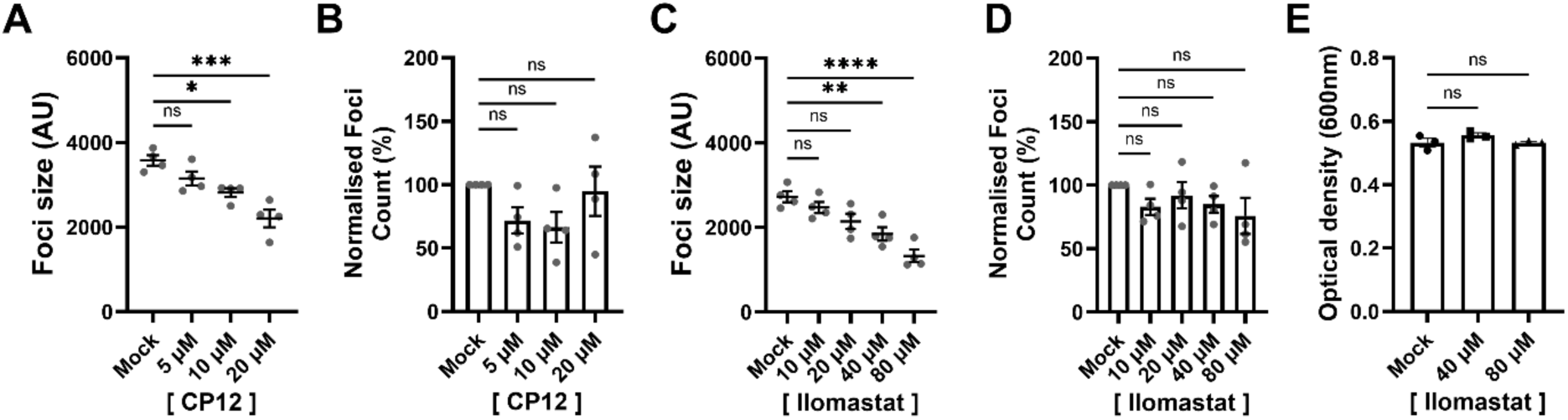
CP12 as a potential matrix metalloproteinase (MMP) inhibitor. (A–B) Quantification of *Shigella* infection focus size (A), expressed in arbitrary units, and number of infection foci (B), normalized to the Mock condition, at 8 h post-infection in the presence of increasing concentrations of CP12. (C–D) Quantification of *Shigella* infection focus size (C), expressed in arbitrary units, and number of infection foci (D), normalized to the Mock condition, at 8 h post-infection in the presence of increasing concentrations of Ilomastat, a broad-spectrum MMP inhibitor. (E) Growth of *Shigella flexneri* in LB medium in the presence of Ilomastat, measured by optical density at 600 nm (OD₆₀₀) after 3 h of incubation. Each dot represents an independent biological replicate. Error bars indicate the standard error of the mean (SEM). Statistical significance was determined by one-way ANOVA followed by Dunnett’s multiple-comparisons test. ns, not significant; **, p < 0.01; ****, p < 0.0001.

### Thiosangivamycin (CP20) inhibits *S. flexneri* invasion and dissemination through CDK2-dependent and CDK2-independent pathways, respectively

We identified thiosangivamycin (CP20, NSC105827 or FLIX1) as one of four compounds that significantly reduced infection foci size in HT-29 monolayers (Supplementary tables 1 and 2). Treatment with thiosangivamycin led to a dose-dependent decrease in *S. flexneri* infection foci size and count (Fig 4A, 4D). As previously described, this compound did not significantly affect ATP levels, as judged by CellTiter-Glo assay, following 8 h of treatment, corresponding to the duration of the infection assay (Supplementary Table 3, Table 1). Although CP20 induced a slight increase in the proportion of nuclei that stained positive by TUNEL assay compared to DMSO alone, it did not trigger HT-29 cell death after 24 h of treatment (Table 1; Supplementary table 3). In addition, it did not exhibit antibacterial activity against *S. flexneri* (Fig. 2A, Fig 2B). Interestingly, a previous study identified this compound as an inhibitor of the Cyclin A2-CDK2 complex ^35^. CDK2 is a kinase involved in cell cycle control ^36^. To confirm that thiosangivamycin inhibits CDK2 and therefore affects the cell cycle, we quantified the number of HT-29 cells at 24h, 48h, and 72h with different concentrations of CP20. As expected, we observed a stabilization of the nuclei count over time with no increase after 72h of incubation (Supplementary Fig 4A, 4B). To confirm that the activity of thiosangivamycin is attributable to CDK2 inhibition, we tested two specific CDK2 inhibitors, INX-315 and CVT-313, that target the ATP binding pocket of the kinase ^37,38^. We first showed that both compounds inhibit HT-29 cell growth in a dose-dependent manner (Supplementary Fig 4C-F). Next, we showed that inhibition of CDK2 by INX-315 and CVT-313 decreases the number of *S. flexneri* foci in a dose-dependent manner, suggesting a role for CDK2 in epithelial cell invasion (Fig 4E-F). However, unlike thiosangivamycin, neither compound affected *S. flexneri* dissemination, suggesting that thiosangivamycin targets additional cellular factors beyond CDK2 that contribute to the reduction in infection foci. (Fig 4B-C). To verify the role of CDK2 inhibition in *S. flexneri* invasion, we added inhibitors following 1 hour of infection, allowing *S. flexneri* to enter HT-29 cells, before adding the inhibitors and killing extracellular bacteria with gentamicin. As expected, foci size and foci number were not affected in the presence of CVT-313 and INX-315, suggesting that CDK2 is important for *S. flexneri* invasion (Fig 4H, 4I, 4K, 4L). Interestingly, thiosangivamycin still presented with a slight decrease in foci size, suggesting that thiosangivamycin inhibits *S. flexneri* dissemination in a CDK2-independent manner (Fig 4G, 4J). To rule out the possibility that impaired invasion resulted from antimicrobial activity of the two CDK2 inhibitors against *S. flexneri*, we tested the two highest concentrations used in our assay for inhibition of bacterial growth in LB broth. Importantly, neither compound affected bacterial growth compared with control cultures, indicating that CDK2 activity contributes to *S. flexneri* invasion of HT-29 epithelial cells without directly influencing bacterial replication (Supplementary Fig 4G, 4H).

**Figure 4:**
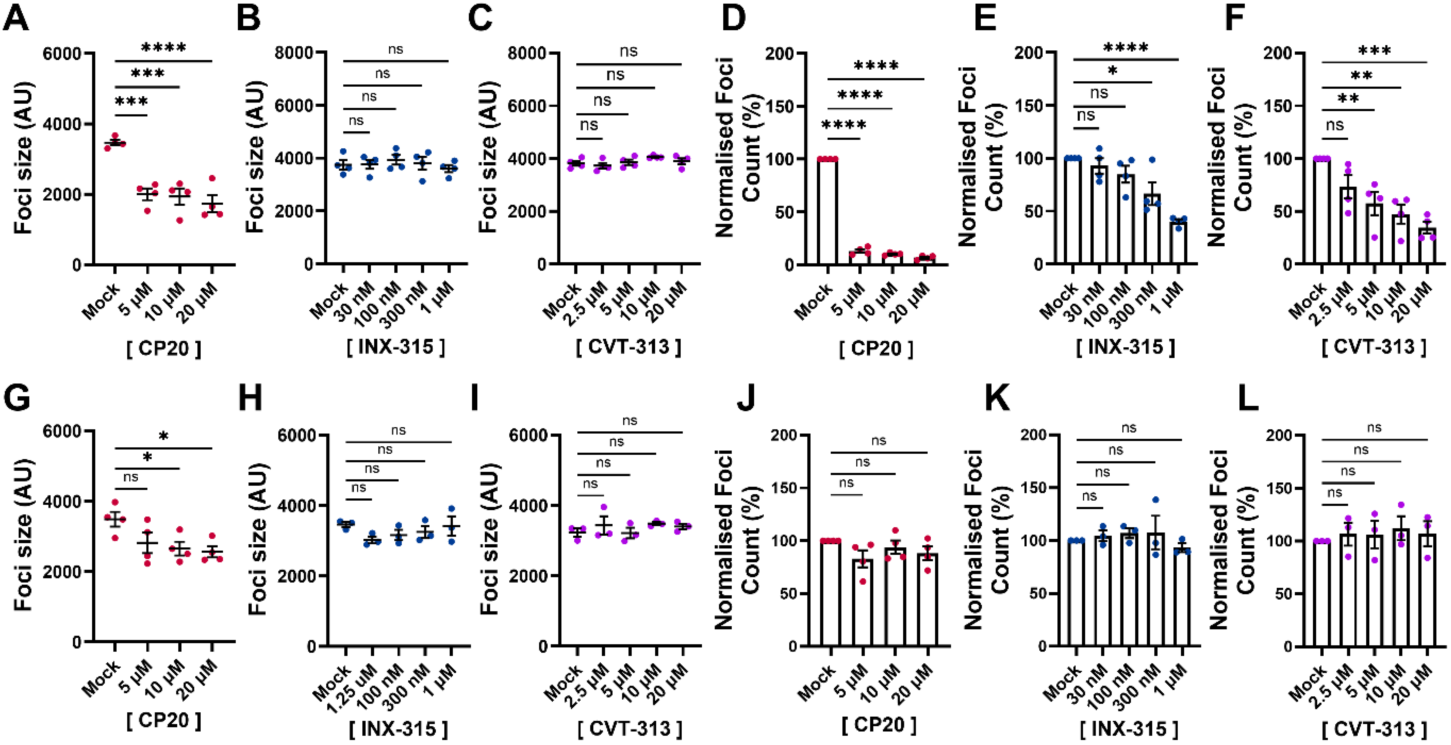
Thiosangivamycin (CP20) inhibits *Shigella* invasion of HT-29 cells through a mechanism recapitulated by CDK2 inhibitors, while inhibiting *Shigella* dissemination through a CDK2-independent pathway. (A–C) Quantification of *Shigella* infection focus size, expressed in arbitrary units, at 8 h post-infection following treatment with increasing concentrations of CP20 (A), INX-315 (B), or CVT-313 (C). Compounds were added 1 h prior to infection. (D–F) Quantification of the number of *Shigella* infection foci, normalized to the Mock condition, at 8 h post-infection following treatment with increasing concentrations of CP20 (D), INX-315 (E), or CVT-313 (F). Compounds were added 1 h prior to infection. (G–I) Quantification of *Shigella* infection focus size, expressed in arbitrary units, at 8 h post-infection following treatment with increasing concentrations of CP20 (G), INX-315 (H), or CVT-313 (I). Compounds were added 1 h after infection. (J–L) Quantification of the number of *Shigella* infection foci, normalized to the Mock condition, at 8 h post-infection following treatment with increasing concentrations of CP20 (J), INX-315 (K), or CVT-313 (L). Compounds were added 1 h after infection. Each dot represents an independent biological replicate. Error bars indicate the standard error of the mean (SEM). Statistical significance was determined by one-way ANOVA followed by Dunnett’s multiple-comparisons test. ns, not significant; *, p < 0.05; **, p < 0.01; ***, p < 0.001; ****, p < 0.0001.

### CP24 (NSC70799) reduces *S. flexneri* infection foci independently of neurosteroid and glucocorticoid signaling

We identified NSC70799 (CP24) as one of four compounds that significantly reduced infection foci size in HT-29 monolayers (Supplementary tables 1 and 2). Treatment with NSC70799 decreased *S. flexneri* infection foci size at 20 µM but not at lower doses (Supplementary Fig 5A). It did not affect bacterial invasion (Supplementary Fig 5E). CP24 did not affect ATP levels following either 8 h of treatment, corresponding to the duration of the infection assay, or following a longer incubation period (Table 1; Supplementary table 3). In addition, it did not exhibit direct antibacterial activity against *S. flexneri* (Fig. 2A, Fig 2B). NSC70799 is steroid-like compound, identified as 11-Hydroxy-11-methylpregnane-3,20-dione dioxime. We tested whether the CP24 inhibitory activity on foci size formation was mediated by neurosteroid or glucocorticoid signaling pathways. To assess whether NSC70799 acts through neurosteroid-like modulation of GABAergic signaling, cells were treated with the GABA(A) receptor agonist gamma-aminobutyric acid (GABA) and the antagonist picrotoxin ^39^. Neither activation nor inhibition of GABA(A) receptors affected *S. flexneri* foci number or size, suggesting that GABAergic signaling does not contribute to bacterial dissemination in HT-29 cells (Supplementary Figs 5B, 5C, 5F, 5G). We tested the potential involvement of the glucocorticoid-signaling pathway using the potent agonist dexamethasone ^40^. Dexamethasone treatment did not recapitulate the NSC70799 phenotype, indicating that the glucocorticoid-signaling pathway does not support *S. flexneri* dissemination in HT-29 cells (Supplementary Figures 5D, 5H). These results suggest that, despite its steroid-like structure, NSC70799 does not exert its inhibitory activity through the canonical neurosteroid or glucocorticoid receptor signaling pathways.

### The adenine analog CP01 (NSC45641) restricts *S. flexneri* dissemination through reversible purine limitation

CP01 (NSC45641) exhibits conditional antimicrobial activity in a nutrient-limited medium, but not in rich medium indicating that its inhibitory activity is dependent on nutrient availability (Fig 2A, 2B). CP01 did not affect HT-29 viability at concentrations up to 100 µM following 24h of incubation (Table 1). In infected HT-29 cell monolayers, CP01 reduced infection focus size in a dose-dependent manner, with statistically significant inhibition observed at concentrations of 10 and 20 µM (Fig. 5A). Infection focus number was reduced by approximately 30% without clear dose dependence (Fig 5B). CP01 (2-Propanone, 2-(9H-purin-6-yl)hydrazone)) is a modified adenine-like derivative containing a purine scaffold, suggesting a potential interaction with purine metabolism (Fig 5C). To characterize its activity under nutrient-limiting conditions, we monitored bacterial growth in minimal medium with increasing concentrations of CP01. CP01 delayed bacterial growth starting at a concentration of 2.5 µM and higher (Fig 5D). Given the *S. flexneri* dissemination defect observed with CP01 treatment, we next tested whether CP01 restricts intracellular bacterial replication. Intracellular CFUs recovered at 6 hours post-infection, following 4 h of compound exposure, was unaffected at concentrations up to 20 µM (Fig. 5E). Although CP01 limited *S. flexneri* growth in nutrient-limited medium, it did not measurably impair intracellular replication, indicating that the reduction in infection focus size is unlikely to result simply from impaired intracellular bacterial growth. In bacteria, adenine, guanine, and hypoxanthine are interconnected through the purine salvage pathway (Fig. 5F). Hypoxanthine is converted into inosine monophosphate (IMP), a central purine intermediate that serves as a precursor for AMP and GMP. Adenine is salvaged directly into AMP, whereas guanine is converted into GMP, thereby supporting nucleotide biosynthesis and cellular growth (Fig. 5F) ^41^. To determine whether CP01 interferes with purine metabolism, we tested whether the bacterial growth defect observed in minimal medium was restored by supplementing the medium with 100 µM adenine, guanosine or hypoxanthine in the presence of 20 µM of CP01. Guanosine was used as a substitute for guanine, because of its higher solubility. Uridine was included as a pyrimidine control. In the absence of CP01, none of these metabolites affected growth (Fig 5G). In the presence of CP01, adenine, hypoxanthine and guanosine restored growth, with adenine providing the most efficient rescue (Fig 5G). Uridine produced a small but detectable effect. The ability of several distinct purine sources to rescue the growth defect indicates that CP01 imposes a global purine limitation rather than blocking a single salvage branch. However, the efficacy of adenine may reflect a difference in salvage pathway efficiency among these substrates.

**Figure 5.**
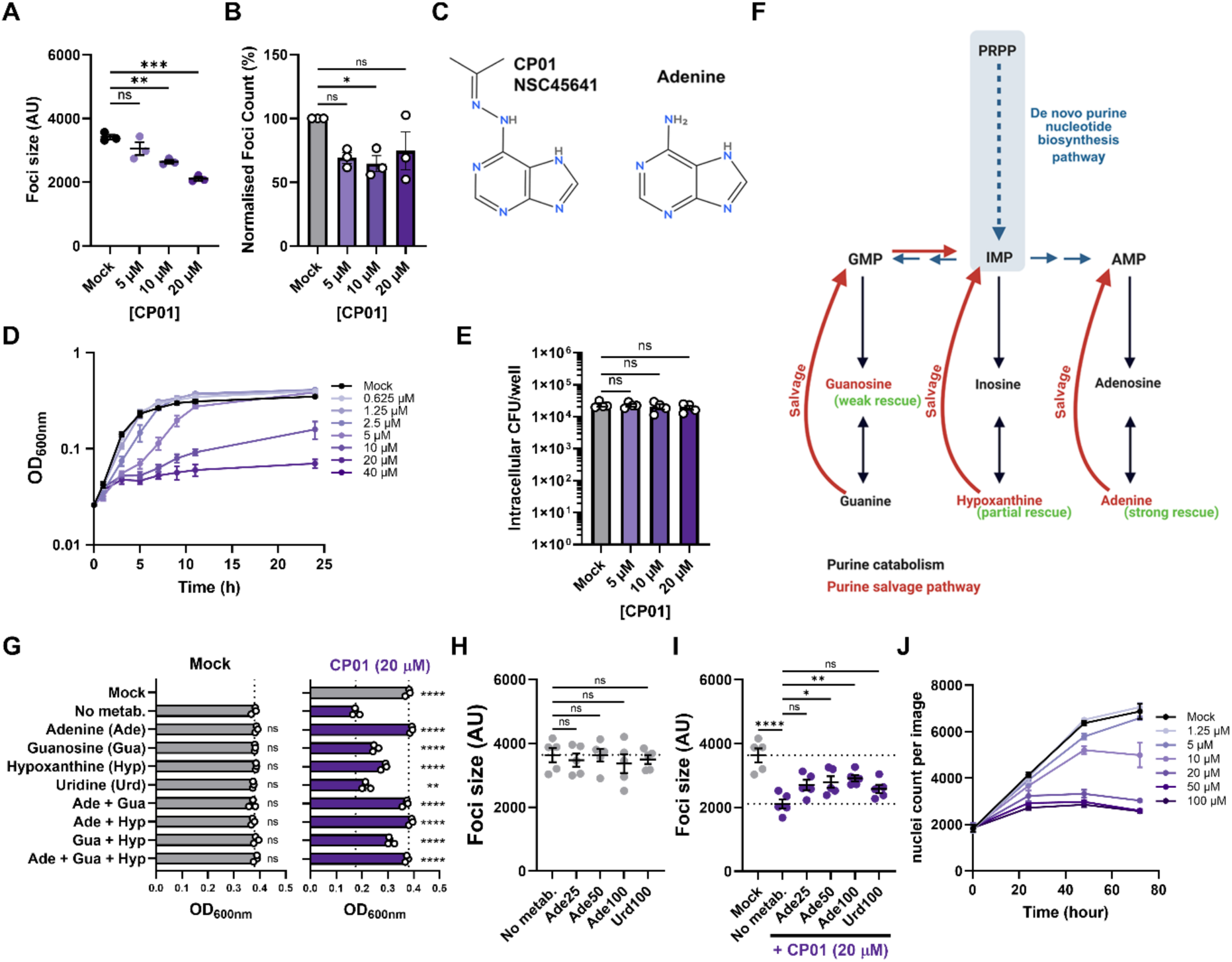
CP01 (NSC45641) restricts Shigella dissemination through reversible purine limitation. (A–B) *Shigella flexneri* 2457T infection focus area (A) and normalized focus number (B) at 8 hpi with increasing CP01 concentrations. (C) Chemical structures of CP01 and adenine. (D) Bacterial growth in minimal medium with increasing CP01 concentrations. (E) Intracellular CFU after 6 h of infection, including 4 h of CP01 treatment. (F) Purine salvage (red), catabolism (black), and de novo biosynthesis (blue) pathways. (G) Growth rescue by adenine (Ade), guanosine (Gua), hypoxanthine (Hyp), or combinations (100 µM), with or without 20 µM CP01; uridine (Urd) served as a negative control. (H–I) Infection focus area in HT-29 monolayers at 8 hpi following adenine supplementation (25–100 µM), without (H) or with (I) 20 µM CP01. (J) HT-29 cell growth following CP01 treatment, measured by nuclei counts over 3 days. Curves represent means of three independent experiments; error bars indicate SEM. Statistical significance was determined by one-way ANOVA with Dunnett’s test. ns, not significant; *, p < 0.05; **, p < 0.01; ***, p < 0.001; ****, p < 0.0001.

We next asked whether adenine supplementation could also restore bacterial dissemination during infection. Adenine and uridine had no effect on infection focus size in the absence of CP01 (Fig 5H). In the presence of 20 µM CP01, adenine partially restored focus size in a dose-dependent manner, whereas uridine did not (Fig 5I), indicating that the purine limitation imposed by CP01 contributes to the dissemination defect in host cells. Finally, because purine metabolic pathways are conserved across organisms, we examined the effect of CP01 on actively dividing HT-29 cells over three days. Proliferation was unaffected at 1.25 µM and only modestly reduced at 5 µM, whereas concentrations of 10 µM and above arrested HT-29 proliferation, with nuclei counts plateauing shortly after treatment and remaining stable for up to 72 hours (Fig 5J). The absence of any decline in nuclei number at concentrations up to 100 µM is consistent with the lack of measurable cytotoxicity on the low-proliferative monolayer (Table 1), indicating that CP01 arrests division of proliferating host cells without killing them.

Together, these results link the conditional antimicrobial activity of CP01 with purine limitation as evidenced by the restoration of bacterial growth with adenine supplementation under both nutrient-restricted conditions and during infection of HT-29 epithelial monolayers. CP01 also inhibits the proliferation of dividing host cells at concentrations comparable to those required for its antibacterial activity.

### CP03, an uracil analog, imposes a pyrimidine limitation on *S. flexneri* during growth and dissemination

CP03 (NSC367428) is a methyl 5-fluoro-4-methoxy-2,6-dioxohexahydro-5-pyrimidinecarboxylate containing a fluorinated pyrimidine scaffold structurally related to uracil and 5-fluorouracil (5-FU), suggesting potential interference with pyrimidine metabolism (Fig. 6A). CP03 was one of the two compounds presenting a conditional inhibition in nutrient-limited medium. To characterize its activity, we monitored bacterial growth in minimal medium with increasing concentrations of CP03. CP03 altered bacterial growth kinetics in a concentration-dependent manner (Fig 6B). As 5-FU is related structurally, we also monitored the effect of 5-FU on *S. flexneri* growth and observed a dose dependent inhibition (Fig 6C). However, 5FU was more potent, inhibiting growth at nanomolar concentrations, below the concentrations required for CP03.

**Figure 6:**
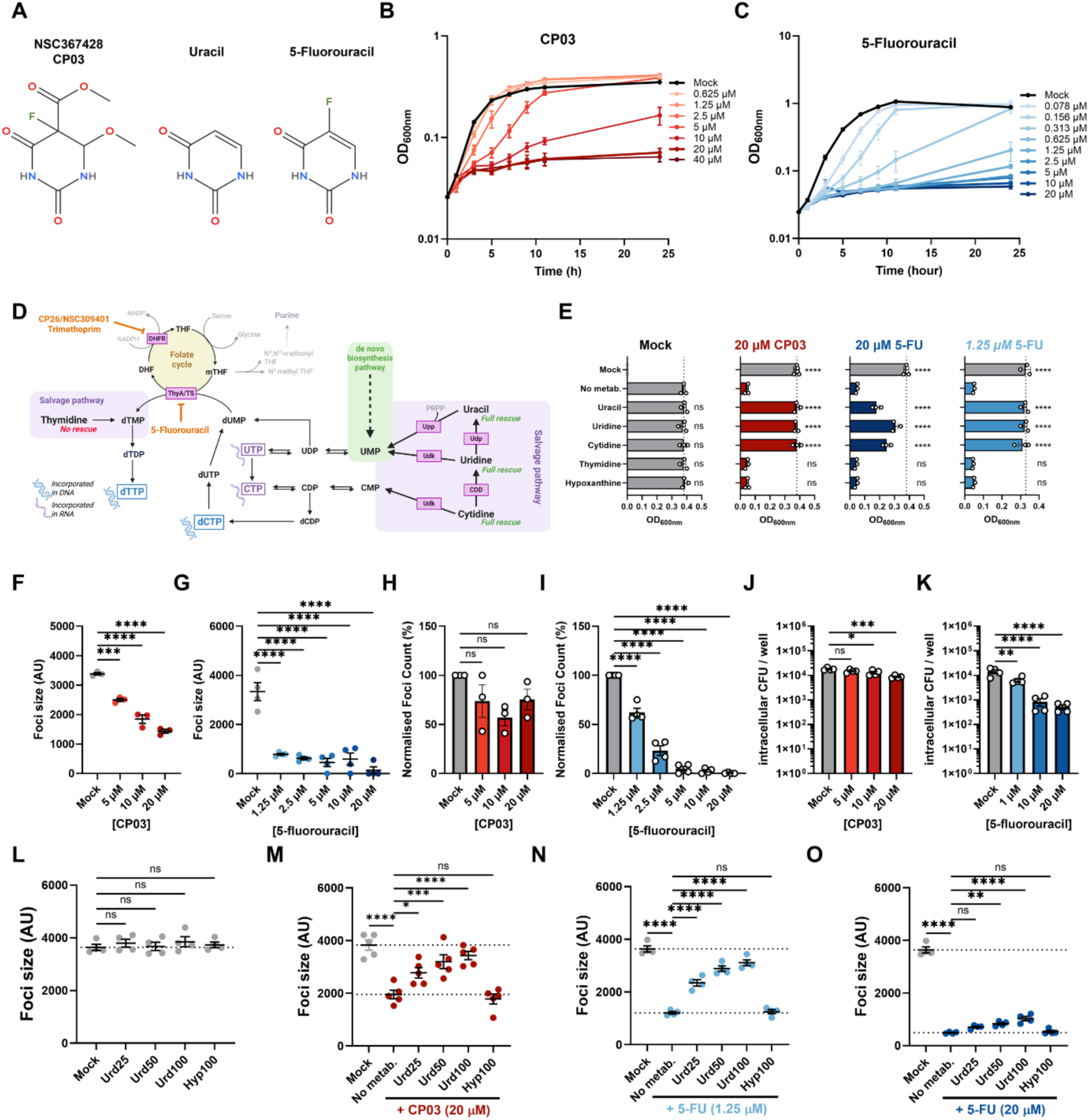
CP03, a uracil-related compound, and its effects on pyrimidine metabolic pathways. (A) Chemical structures of CP03, a uracil-related compound, uracil, and 5-fluorouracil (5-FU). (B–C) Growth curves of *Shigella flexneri* in minimal medium in the presence of increasing concentrations of CP03 (B) or 5-FU (C). (D) Simplified schematic representation of pyrimidine metabolism, highlighting the de novo biosynthesis and salvage pathways of uridine, uracil, cytidine, and thymidine. Pyrimidine nucleotides are synthesized through the de novo pathway (green), which generates UMP as a central intermediate, or through salvage pathways involving the recycling of nucleobases and nucleosides. Uracil is converted into UMP by uracil phosphoribosyltransferase (Upp), while uridine is converted into UMP by uridine kinase (Udk) or interconverted with uracil by uridine phosphorylase (Udp). Cytidine can be salvaged through Udk-mediated phosphorylation to CMP or through cytidine deaminase (Cdd). UMP is subsequently phosphorylated and interconverted through the UDP/UTP and CDP/CTP nucleotide pools, which contribute to RNA synthesis (purple single-stranded helix). Following ribonucleotide reduction, dCTP and dTTP are incorporated into DNA (blue double-stranded helix). dTMP is generated from dUMP by thymidylate synthase (ThyA/TS), a reaction linked to the folate cycle (yellow), or through the thymidine salvage pathway (purple). The 5-FU metabolite FdUMP and trimethoprim are known inhibitors of the folate-dependent pathway, acting on ThyA/TS and dihydrofolate reductase (DHFR), respectively. CP26/NSC309401, identified in this screen, has previously been reported to inhibit DHFR. Metabolites tested for their ability to rescue bacterial growth are highlighted in green (complete rescue) or red (no rescue). (E) Bacterial growth rescue experiments in minimal medium following supplementation with 100 µM uracil, uridine, cytidine, or thymidine in the presence or absence of 20 µM CP03 (red), 20 µM 5-FU (dark blue), or 1.25 µM 5-FU (light blue). Hypoxanthine, a purine metabolite, was used as a negative control. (F–G) Quantification of *Shigella flexneri* 2457T infection focus area in HT-29 cell monolayers at 8 h post-infection (hpi) in the presence of increasing concentrations of CP03 (F) or 5-FU (G). (H–I) Quantification of the number of *Shigella flexneri* 2457T infection foci in HT-29 cell monolayers at 8 hpi, normalized to the Mock condition, following treatment with increasing concentrations of CP03 (H) or 5-FU (I). (J–K) Intracellular CFU recovered after 6 h of infection, including 4 h of incubation in the presence of CP03 (J) or 5-FU (K). (L–O) Quantification of *Shigella* infection focus area in HT-29 cell monolayers at 8 hpi following supplementation with increasing concentrations of uridine (25 µM [Urd25], 50 µM [Urd50], or 100 µM [Urd100]) in the absence of treatment (L) or in the presence of 20 µM CP03 (M), 1.25 µM 5-FU (N), or 20 µM 5-FU (O). Hypoxanthine (100 µM; Hyp100) was used as a negative control. Compounds and metabolites were added at 1 hpi. Each dot represents an independent biological replicate. Error bars indicate the standard error of the mean (SEM). Statistical significance was determined by one-way ANOVA followed by Dunnett’s multiple-comparisons test. ns, not significant; **, p < 0.01; ***, p < 0.001; ****, p < 0.0001.

In bacteria, uracil and uridine are salvaged into uridine monophosphate (UMP), a central precursor for pyrimidine nucleotide biosynthesis. UMP can subsequently be converted into UDP, UTP, and CTP, thereby supporting RNA synthesis among many cellular processes ^41^. Cytidine can also contribute to pyrimidine nucleotide biosynthesis through conversion into CMP and subsequently CTP, or by deamination into uridine by cytidine deaminase (Cdd). Thymidine is phosphorylated into dTMP by thymidine kinase, which replenishes the dTMP/dTTP pool required for DNA synthesis (Fig. 6D). To determine whether CP03 and 5-FU interfere with pyrimidine metabolic pathways, we tested whether supplementing the medium with 100 µM uridine, uracil, cytidine, thymidine or the purine metabolite hypoxanthine as negative control, would rescue bacterial growth. In the absence of compounds, none of these metabolites altered growth. Uracil, uridine and cytidine restored *S. flexneri* growth in the presence of 20 µM CP03, but thymidine and hypoxanthine did not (Fig. 6E). A similar pattern was observed for 5-FU at 1.25 µM and 20 µM, but growth restoration was incomplete at the higher dose. These findings indicate that CP03 affects pyrimidine metabolism at or upstream of the uridineand cytidine-associated nucleotide pools rather than limiting dTMP availability. CP03 inhibited dissemination in HT-29 monolayers in a dose-dependent manner (Fig 6F), whereas a significant reduction in infection foci number was observed only at the highest concentration tested (Fig 6H). In contrast, 5-FU inhibitory effect on both infection focus size and number was more pronounced, with the number of detectable foci markedly reduced at concentrations of 5 µM and above (Fig 6G, 6I), consistent with its greater potency in minimal medium and with its antimicrobial activity previously reported in *E. coli* ^42^. We next investigated whether these compounds restricted intracellular bacterial replication. Intracellular CFUs recovered at 6 hpi, following 4 h of compound exposure, were modestly but significantly reduced by CP03 at 10 and 20 µM (Fig 6J), whereas 5-FU led to a greater than tenfold reduction at 20 µM (Fig 6K). CP03 therefore restricts intracellular replication in addition to limiting dissemination, distinguishing its activity from that of CP01. We then tested whether uridine supplementation could restore bacterial dissemination during infection of an HT-29 cell monolayer. Uridine alone had no effect on infection focus size (Fig 6L). In the presence of 20 µM CP03 added 1 h after infection, uridine restored focus size in a dose-dependent manner, with nearcomplete recovery at 100 µM, whereas hypoxanthine had no detectable effect (Fig 6M). Similarly, uridine rescued the inhibitory effect of 5-FU at 1.25 µM (Fig 6N) but provided only a slight recovery at 20 µM 5-FU, a concentration at which dissemination was almost completely abolished (Fig. 6O).

Together, these results indicate that CP03 imposes a pyrimidine-associated metabolic constraint that can be bypassed through uracil, uridine or cytidine salvage. This links CP03 conditional antimicrobial activity to the disruption of pyrimidine nucleotide homeostasis, impacting both bacterial growth and intracellular dissemination.

### Upp-dependent activation is required for CP03 and 5-FU activities

5-FU is a fluorinated derivative of uracil that is metabolized by bacteria, leading to the formation of FdUMP and FUTP (Fig 7A). FdUMP inhibits thymidylate synthase (ThyA/TS) by occupying the active site of the enzyme, which results in blocking dTMP formation. This leads to severe depletion of the dTTP pool and block in DNA synthesis ^42,43^. FUTP is structurally close to UTP and can be incorporated in RNA by the RNA polymerase, which disrupts normal RNA function and processing, ultimately impairing protein synthesis. Given their structural similarities and comparable effect on *S. flexneri* growth in minimal medium and dissemination in HT-29, we hypothesized that CP03 could exert its antimicrobial activity through a mechanism similar to that of 5-FU. Conversion of 5-FU to 5-fluoro-UMP in bacteria requires the presence of the uracil phosphoribosyltransferase enzyme (Upp). To test the requirement of Upp in 5-FU conversion, we generated a *S. flexneri* Δ*upp* mutant and a complemented strain expressing *upp* from an arabinose-inducible promoter (Δ*upp*-compl). In minimal medium, wild-type *S. flexneri* was inhibited by both CP03 and 5-FU, whereas the Δ*upp* mutant and the complemented strain grew similarly to the mock condition in absence of arabinose (Fig 7B). Induction of *upp* with addition of 0.5% arabinose fully restored sensitivity to both compounds, confirming that Upp is necessary and sufficient for their activity (Fig 7B).

**Figure 7.**
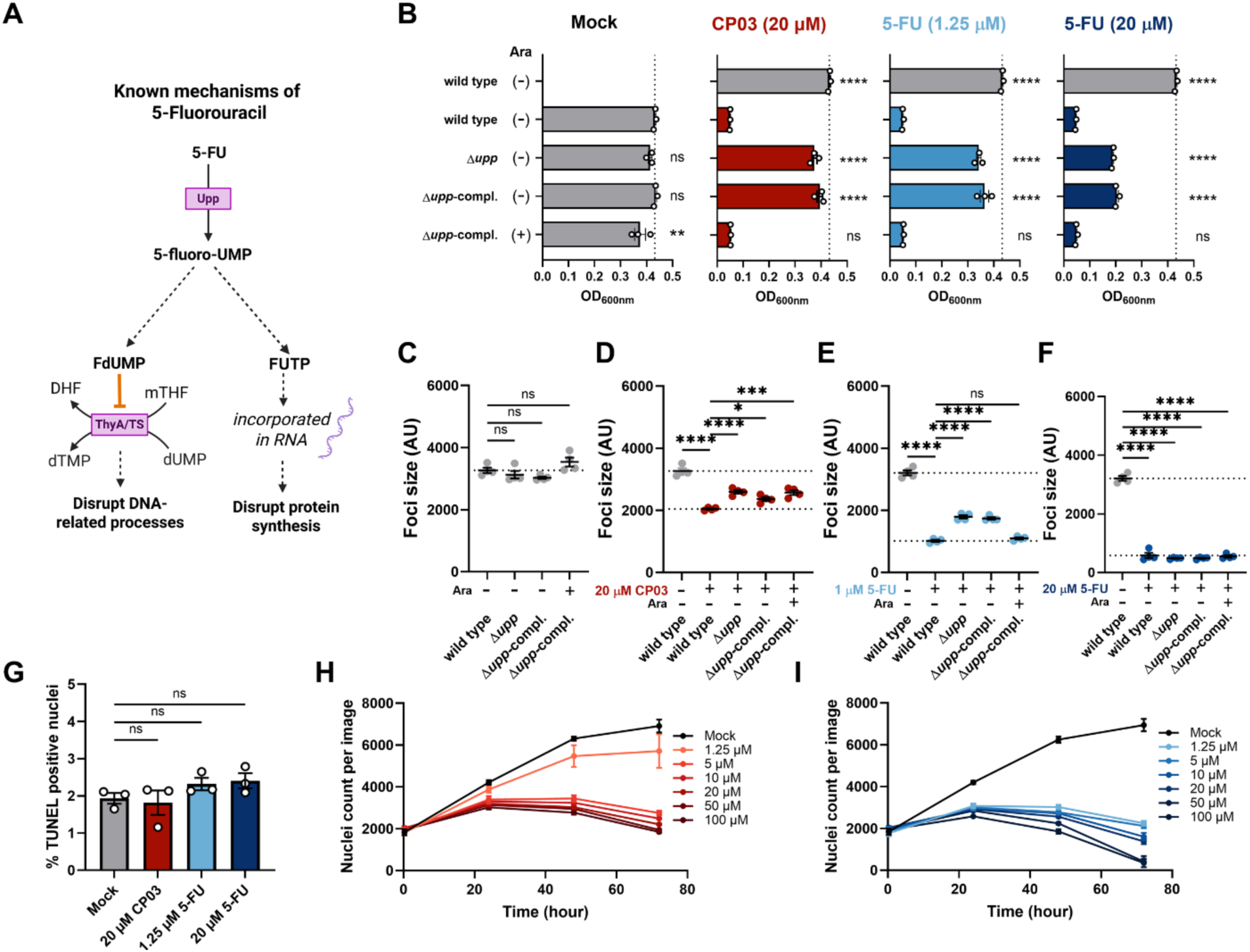
Upp-dependent activation is required for the antibacterial activities of CP03 and 5-FU. (A) Schematic of 5-FU activation by bacterial Upp to generate 5-fluoro-UMP, which is further metabolized via the pyrimidine pathways in Fig. 6D to produce FdUMP and FUTP, disrupting DNA metabolism and RNA synthesis, respectively. (B) OD₆₀₀ of wild-type *Shigella flexneri*, Δupp, and Δupp-compl after 5 h of growth in minimal medium containing 20 µM CP03, 1.25 µM 5-FU, or 20 µM 5-FU. *upp* expression in the complemented strain was induced with 0.5% arabinose, as indicated. (C–F) Infection focus area in HT-29 monolayers at 8 h postinfection with wild-type *S. flexneri*, Δupp, or Δupp-compl, without treatment (C) or following treatment with 20 µM CP03 (D), 1.25 µM 5-FU (E), or 20 µM 5-FU (F). Arabinose induced *upp* expression in the complemented strain. (G) Percentage of TUNEL-positive cells after 24 h of CP03 or 5-FU treatment. (H–I) HT-29 cell growth following treatment with increasing concentrations of CP03 (H) or 5-FU (I), measured by nuclei counts per image over 3 days. Curves show means of three independent experiments. Dots represent independent biological replicates; error bars indicate SEM. Statistical significance was assessed by one-way ANOVA with Dunnett’s multiple-comparisons test. ns, not significant; **, p < 0.01; ***, p < 0.001; ****, p < 0.0001.

We next tested the impact of Upp on compound-mediated growth inhibition during infection of HT-29 monolayers. Without compounds, all strains formed similarly sized infection foci, indicating that loss of *upp* does not affect dissemination (Fig. 7C). At 1.25 µM 5-FU, both the Δ*upp* mutant and complemented strain showed partial but reproducible restoration of infection focus size, which was abolished by arabinose induction of the complemented strain (Fig. 7E), recapitulating results in minimal medium. In contrast, loss of *upp* conferred only modest recovery from CP03-mediated inhibition, with no discernible difference between induced and uninduced complemented strains (Fig. 7D). Thus, a substantial component of CP03-mediated inhibition of bacterial dissemination appears independent of Upp activity. At 20 µM 5FU, dissemination was abolished in all strains, regardless of *upp* genotype (Fig. 7F). This residual Upp-independent activity may reflect effects of CP03 and 5-FU on host-cell physiology. Neither compound reduced viability or increased TUNEL-positive nuclei in 4-day-old HT-29 monolayers after 24 h of treatment (Table 1; Fig. 7G). However, both inhibited HT-29 cell proliferation. CP03 partially inhibited proliferation at 1.25 µM and completely blocked it at 5 µM and above, with nuclei counts plateauing through 72 h (Fig. 7H). 5-FU also arrested proliferation, but nuclei counts fell below baseline across the tested concentration range, indicating cell loss rather than arrest alone (Fig. 7I). Thus, although both compounds require Upp for antibacterial activity, CP03 has a milder effect on host cells, blocking cell division without causing cell loss.

## Discussion

The rapid emergence of MDR and XDR *S. flexneri* strains together with the lack of universal licensed vaccines highlights the need for alternative therapeutic strategies that complement the use of conventional antibiotics. Because epithelial dissemination is central to *S. flexneri* pathogenesis and tissue destruction, targeting host pathways required for cellto-cell spread represents an attractive anti-virulence strategy. In this study, we performed a high-content microscopy-based phenotypic screen of the NCI Diversity Set VII for small molecules that inhibit *S. flexneri* cell-to-cell spread in epithelial monolayers. By counter-screening the primary hits, we identified six structurally diverse compounds that reduced infection foci size without affecting host cell viability in the confluent monolayer. Four compounds (CP08, CP12, thiosangivamycin and CP24) presented no direct effect on bacterial growth supporting a host-directed mechanism of action. The remaining two (CP01 and CP03) displayed antibacterial activity only under nutrient-limited conditions, which conventional antimicrobial screening conditions do not sample. CP03 also reduced intracellular replication in HT-29 cells. Most screening media are enriched in nutrients and metabolites, such as purines, pyrimidines, amino-acids and peptides, and therefore do not reflect the physiological constraints encountered by pathogens within the host environment. We acknowledge that our assay focuses on the epithelial phase of *S. flexneri* infection, using HT-29 cells in absence of other host components, such as immune cells, mucus layer, microbiota and the mechanical forces exerted by peristalsis. Consequently, compounds acting specifically on these host factors may not be identified using our screening approach.

### Compounds with direct activity against *S. flexneri*

As anticipated for a phenotypic screen scoring infection foci size formed by *Shigella* in HT-29, any compounds presenting anti-bacterial activities would lead to a decrease or abolition of the infection. In our screen, eleven of the nineteen validated compounds inhibited *S. flexneri* growth in rich medium. Several of these compounds were already well-described antimicrobials such as Nifuratrone (CP02/NSC303603), bactobolin (CP29/NSC325014) or Streptovaricin C (CP34/NSC199990) ^44–46^. Other identified hits have previously been reported to target essential bacterial enzymes or pathways. CP26 (NSC309401) targets DHFR in *E. coli* ^47^. CP30 (NSC65238) inhibits two-component systems in *Salmonella* ^48^. CP16 (NSC30205) inhibits the highly conserved polyphosphate kinase 2 in *Mycobacterium tuberculosis* without being cytotoxic ^49^. Adenosine 5’-sulfamate (CP21/NSC133114) was described to show antimalarial activity by targeting aminoacyl tRNA synthetases ^50^. Other identified hits were reported as anti-cancer. CP05 (NSC59984) and CP09 (XI-006 or NSC207895) act on the p53 pathway ^28,51^. CP32 (NSC277184) sensitizes cancer stem cells to apoptosis ^52^. However, CP22 (NSC339316) was not found in any study to our knowledge. Four of the hits (bactobolin, CP09, CP26 and Streptovaricin C) were characterized as anti-microbial in a study based on screening the NCI diversity set V for anti-methicillin-resistant *Staphylococcus aureus* (MRSA) ^53^. We note that NSC343256 (hitachimycin) and three compounds: NSC8675 (ethyl violet), NSC345647 (chaetochromin) and NSC70931 (celastrol), were hits in both the antiMRSA compounds study and in our initial screening (Supplementary. Table 1), suggesting a broad spectrum activity spanning Gram-positive and Gram-negative species. These compounds likely acted as conventional antibiotics, and were not further characterized in this study.

### Identifying new factors and pathways through this screen

Cell-to-cell spread drives the epithelial fenestration and tissue destruction that characterize bacillary dysentery, and *S. flexneri* mutants defective in cell-to-cell spread are markedly attenuated *in vivo* ^16,20,21^. We previously showed that inhibition of the host lipid kinase PIK3C3/hVPS34 with compound SAR405 reduces dissemination in epithelial monolayers and colonization of the colonic mucosa in the infant rabbit model ^17^. In the present study, SAR405 was included as a positive control on every screening plate enabling reliable calibration across the screen. The four compounds identified and validated in our study, CP08, CP12, thiosangivamycin and CP24, reduced infection foci size without affecting bacterial growth in either rich or nutrient-limited medium, consistent with a host-directed mode of action. These compounds therefore represent lead candidates whose chemical scaffolds could be further optimized to improve potency. Established animal models of shigellosis such as the infant rabbit model will be a useful pre-clinical platform for evaluating their *in vivo* efficacy.

Ilomastat, a broad-spectrum metalloprotease inhibitor, reduced infection focus size in a dose-dependent manner without affecting focus number or bacterial growth, implicating host metalloprotease activity in cell-to-cell spread. CP12 elicited a similar phenotype and is structurally identical to a compound disclosed in WO2008063667, a patent covering thieno-fused pyrimidin-4-ones as inhibitors of MMP-13, MMP-8, MMP-3 and MMP-2 ^33^. CP12 lacks the zinc-binding group characteristic of firstand second-generation metalloprotease inhibitors. However, inhibitors lacking this group can act through interaction with the S1 pocket rather than the catalytic zinc ion ^54^. Whether CP12 and Ilomastat act on a common target remains to be determined. To our knowledge, no secreted zinc metalloprotease has been characterized as a *S. flexneri* virulence factor. Nevertheless, *S. flexneri* preferentially targets tricellular junctions to spread into adjacent cells and can modulate cell junction through the T3SS bacterial effector IpaC ^55,56^. Given the established role of matrix metalloprotease in remodelling cell junctions and extracellular matrixes, their potential role in *S. flexneri* pathogenesis warrants further investigation ^57,58^. These observation raise the possibility that host MMP activity contribute to the cellular remodeling required for efficient cell-to-cell spread.

Thiosangivamycin (CP20) displayed a more informative phenotype. Its ability to decrease infection focus number was recapitulated by two selective CDK2 inhibitors targeting the ATP pocket, neither of which affected bacterial growth. This phenotype was lost for any of the three compounds when added after the invasion step, supporting a role for CDK2 during *S. flexneri* entry into epithelial cells. By contrast, the reduction of focus size following thiosangivamycin treatment was not reproduced by CDK2 inhibitors, suggesting that this effect involves another molecular target. Thiosangivamycin is a structural derivative of sangivamycin which has been reported to inhibit Protein Kinase C, Haspin kinase and other enzymes ^59–61^. Thus, the pleiotropic effects of thiosangivamycin may involve several targets overlapping with those of sangivamycin.

CDK2 has been implicated as a host factor in several viruses, contributing to viral RNA synthesis and lytic replication. rather than viral entry ^62,63^. To our knowledge, a requirement for the entry step of a bacterial pathogen has not been reported. Moreover, no CDK2 substrate linking the kinase activity to the actin cytoskeleton or the endocytic machinery has been described. Nevertheless, the role of cell cycle on actin-related processes during infection have been reported for several pathogens. For example, *Campylobacter jejuni* preferentially invades G1-phase cells, while *S. flexneri* effector IpaB targets Mad2L2 to interfere with the host cycle progression ^64,65^. Determining whether CDK2 represent a general or a pathogenspecific requirement for bacterial entry will require comparisons across various entry mechanisms, including T3SS-mediated invasion by *S. flexneri* and *Salmonella,* as well as receptor-mediated entry by *Listeria*. Extending these studies to phagocytic uptake by macrophages would add an additional level of investigation, as differentiated macrophages and nonproliferative cells that do not rely on cell-cycle progression.

For NSC126224 (CP08) and NSC70799 (CP24), we could not identify any specific molecular targets or pathways accounting for the observed phenotypes. NSC126224 has been reported as a TLR2 and TLR4 antagonist in murine macrophages. However, inhibition of TLR4, TLR2, NF-κB or nitric oxide synthase did not reduce infection focus size in our model. This may be consistent with differences in receptor expression between intestinal epithelial cells and myeloid cells ^66^. NSC70799 has a steroid-like structure but its associated phenotype was not phenocopied by neither GABA(A) receptor modulation nor glucocorticoid receptor agonism. Importantly, target engagement by the corresponding tool inhibitors was not independently verified in HT-29 cells. Consequently, these findings argue against the involvement of the specific pathways tested in promoting bacterial dissemination under our experimental conditions, but do not establish the molecular mechanisms underlying the effects of CP08 or CP24.

### Intracellular metabolic constraints and *S. flexneri* dissemination

The two remaining compounds inhibited bacterial growth, but only under nutrient-limited conditions, a growth regime that conventional antimicrobial screening approaches do not capture. *S. flexneri* invades and replicates in the cytosol of epithelial and non-epithelial cells, where all nutrients are acquired from the host cell ^67–69^. Here, we identified two compounds acting on purine and pyrimidine metabolic pathways that exhibited conditional antibacterial activity in nutrient-limiting medium and inhibited *S. flexneri* dissemination in HT29 cells. Neither compound affected cell viability or induced cell death in 4-day old monolayers. However, both compounds inhibited HT-29 cell proliferation. Purine and pyrimidine *de novo* biosynthesis and salvage pathways play a very important role in bacterial replication, growth and survival and therefore microbial pathogenesis ^70,71^. In another intracellular bacterium, *Listeria monocytogenes*, cytosolic replication depletes the pool of local nutrients so that cell-to-cell spread becomes necessary to sustain growth of the bacterial population ^72^. Accordingly, compounds that impair nucleotide salvage pathways, or increase bacterial reliance on these pathways would be expected to display their activity in the host cytosol while remaining inactive in nutrient-rich broth cultures.

We identified NSC45641 (CP01) as an adenine-related analog whose inhibitory activity on bacterial growth is alleviated when the growth medium is supplemented with adenine, guanosine or hypoxanthine, with adenine presenting the most prominent effect. The ability of several distinct purine sources to rescue growth inhibition suggests that NSC45641 induces a general limitation in purine availability rather than interfering with a single salvage pathway. However, the more pronounced rescuing effect of adenine may reflect differences in salvage efficiency among purine metabolites or structural similarities that could enable adenine to compete with NSC45641 for binding to its molecular target(s). Previous studies showed that adenine auxotrophy only slightly reduced *S. flexneri* virulence *in vivo,* whereas guanine auxotrophy strongly attenuated virulence both *in vivo* and *in vitro* ^73,74^. In fact, the use of mutant in the *guaBA* operon involved in the guanine metabolic pathway were proposed as *S. flexneri*-based vaccine vectors and have provided encouraging results in terms of protection and immunogenicity ^75,76^.

Our screen identified NSC367428 (CP03), a compound with a fluorinated pyrimidine scaffold structurally related to uracil and 5-fluorouracil (5-FU). NSC367428 phenocopies the activity of 5-fluorouracil (5-FU) across all levels investigated. Both compounds inhibit bacterial growth in minimal medium, an effect that can be rescued by uracil, uridine, and cytidine, but not thymidine. Furthermore, their inhibitory effects on intracellular replication and cell-to-cell spread are rescued by uridine, and their activity is dependent on Upp. However, NSC367428 differs from 5-FU in terms of potency, which may reflect differences in membrane permeability, susceptibility to Upp-mediated activation, or binding affinity for their respective molecular targets. The observed Upp dependence is also noteworthy from a therapeutic perspective. Unlike bacteria, mammalian cells lack a dedicated uracil phosphoribosyltransferase and instead activate fluoropyrimidines through orotate phosphoribosyltransferase (OPRT), or through the sequential activities of uridine phosphorylase and uridine kinase ^43^. These distinct activation pathways suggest a potential opportunity for selective antibacterial targeting. However, while NSC367428 and 5-FU do not trigger cell death (TUNEL assay) nor affect cell viability on a low-proliferating HT-29 cell monolayer, both inhibited HT-29 proliferation. This is consistent with the established antiproliferative activity of 5-FU, a chemotherapeutic agent used to treat several cancers and skin conditions ^77,78^. Although short-term fluoropyrimidine treatment could potentially offer an alternative to conventional antibiotics for shigellosis, gastrointestinal toxicity and the impact on epithelial regeneration remain important concerns. Furthermore, direct bacterial targeting raises the risk of resistance development, as illustrated by the emergence of 5-FU and 5-fluorocytosine through loss-of-function mutations in the *upp* gene of *Pseudomonas aeruginosa* ^79^.

Taken together, the two compounds illustrate that the two nucleotide branches are not equivalent constraints for cytosolic *S. flexneri*. Purine limitation by NSC45641 reduced dissemination without a detectable effect on intracellular replication, consistent with the mild phenotype of adenine auxotrophs, whereas pyrimidine limitation by NSC367428 and 5-FU affected both replication and spread. Nucleotide analogs active only under nutrient restriction therefore illustrate a category of conditional antimicrobial that conventional broth-based screening scores as inactive. Whether this category extends beyond nucleotide antimetabolites, and whether it generalises beyond the single chemical library and cell type sampled here, remain to be established.

## Conclusion

In summary, we recovered two classes of compound that conventional antibacterial screening would overlook. Four candidate host-directed inhibitors of cell-to-cell spread, two of which implicate host metalloprotease activity and host CDK2 in distinct steps of the infection cycle, and two nutrient-conditional antimicrobials, for one of which deletion and complementation of *upp* establish a requirement for activation. Individual compounds will require genetic validation of their host targets, chemical characterization and testing in our established infant rabbit model to further assess their therapeutic potential.

## Materials & Methods

### Bacteria and growth culture

The wild type *S. flexneri* strain used in this study was *S. flexneri* 2a str. 2457T ^80^. LB agar plate supplemented with 0.01% Congo Red and LB broth (BP1426-2, Fischer) was used to grow *S. flexneri* strains. *Escherichia coli* strains (DH5α, *λpir and* Δ*nic35*) were grown in LB broth with appropriate antibiotic. E.coli Δ*nic35* strain was grown by supplementing the medium with 100 µg/ml 2,6-diaminopimelic acid (DAP). For bacterial growth assay, *S. flexneri* strains were grown in minimal medium consisting of M9 minimal salts (M9956, Sigma) supplemented with 2 mM of MgSO4 (M7506, Sigma-Aldrich), 0.1 mM CaCl2 (C-5080; Sigma), 0.4 % Glucose (1916-01, J.T.Baker), 12.5 µg/ml Nicotinic acid (N4126, Sigma), 45 µg/ml Methionine (M9625, Sigma) and 20 µg/ml Tryptophan (T0254, Sigma-aldrich) ^25^. When appropriate, medium was supplemented with chloramphenicol (10 µg/mL final concentration), ampicillin (100 µg/mL final concentration), tetracycline (10 µg/mL final concentration), kanamycin (30 µg/mL final concentration) or arabinose (0.5%).

To visualize bacteria by fluorescence microscopy, the pMMB207 plasmid conferring chloramphenicol resistance and harboring the gene encoding cyan fluorescent protein (CFP) under the control of an isopropyl-beta-D-thiogalactopyranoside (IPTG)-inducible promoter was introduced by electroporation, resulting in a CFP-expressing *S. flexneri* strain, as previously described ^21^.

### Generation of *upp* knockout mutant and complemented strain

The Δ*upp* strain of *S. flexneri* was generated by allelic exchange, resulting in replacement of the open reading frame (ORF) of *upp* (SF2542) by the ORF of a kanamycin resistance gene, as previously described for the generation of other mutants ^20,21,81^. Briefly, an overlap PCR construct of the upstream region of *upp*, the coding sequence of kanamycin, and the down-stream region of *upp* was cloned into pSB890-cat suicide vector using the primers listed in supplementary table 4. pSB890-cat carried a tetracycline resistant cassette for selection. The fragment obtained by overlap PCR was digested with NotI and ligated into a NotI/SmaI digested pSB890-cat suicide vector. The ligation product was transformed into the λpir *Escherichia coli* strain to select for clones. The plasmid was verified using whole-plasmid sequencing (Eurofins) and transformed in *E. coli* Δ*nic35*. The *S. flexneri upp::Kan* strain was generated by conjugation with *E. coli Δnic35* harboring the suicide vector pSB890 mentioned earlier. Briefly, 900 µl of a culture of the *E. coli Δnic35* strain harboring the plasmid, grown overnight, was mixed with 300 µl of a culture of the *S. flexneri* 2457T strain grown overnight. The mixture was pelleted via centrifugation, and the dense bacterial pellet was spotted onto an LB agar plate supplemented with DAP. The plate was incubated at 30°C for 24 h. After the incubation period, the spot was collected in sterile phosphate-buffered saline (PBS). Serial dilutions were plated onto LB agar supplemented with kanamycin and incubated at 37°C. Colonies were streaked onto LB agar (without NaCl) supplemented with 10% sterile sucrose (Fisher Bioreagents) and kanamycin. Mutants were selected on these plates after incubation at 30°C for 24 h.

The Δ*upp* mutant was complemented (Δ*upp*-compl) by expressing *upp* from the arabinose-inducible pBAD promoter in vector pBAD18 (ATCC 87393). Primers used for creating the pBAD18-upp vector are listed in supplementary table 4. The *upp* PCR fragment was generated by overlapping PCR and ligated into the pBAD18 vector (ATCC 87393) after digestion with SacI and XbaI restriction enzymes. The clone was transformed into *E. coli DH5α*. The plasmid vector was extracted from the clone, verified by whole-plasmid sequencing, and introduced into the *S. flexneri upp* mutant by transformation. When appropriate, arabinose at 0.5 % final was added to the medium to induce the expression of *upp*. Ampicillin (100 µg/ml final) is used as selective pressure to maintain the plasmid.

### HT-29 culture maintenance

HT-29 cell line was cultured at 37°C with 5% CO2 in McCoy’s 5A medium (Gibco) supplemented with 10% heat-inactivated fetal bovine serum (HI-FBS; Gibco). Dulbecco’s Phosphate Buffered Saline (DPBS; Gibco) was used to wash the cells. HT29 was lifted with Trypsin-EDTA (Gibco) at 0.25 % for 10 minutes.

### Chemical compounds used in this study

PIK3C3 inhibitors SAR405 (20 mM stock in DMSO, Cat#S7682, Selleckchem) was used as control along this study as it was previously shown to inhibit *S. flexneri* cell-to-cell spread resulting in smaller foci ^17^. The other inhibitors used in this study are the TLR4 inhibitor TAK-242 (20 mM stock in DMSO, Cat#614316, Sigma-Aldrich), the TLR2 inhibitor C29 (20 mM stock in DMSO, Cat#HY-100461, MedChemExpress), the NFkB inhibitor BAY 11-7082 (20 mM stock in DMSO, Cat#HY-13453, MedChemExpress), the iNOS inhibitor 1400W dihydrochloride (20 mM stock in water, Cat#HY-18731, MedChemExpress), the NOS inhibitor L-NAME hydrochloride (300 mM stock in water, Cat#HY-18729A, MedChemExpress), the metalloprotease pan-inhibitor Ilomastat (10 mM stock in DMSO, Cat#M5939, Sigma-Aldrich), the CDK2 inhibitor INX-315 (10 mM stock in DMSO, Cat#HY-162001, MedChemExpress), the CDK2 inhibitor CVT-313 (20 mM stock in DMSO, Cat#HY-15339, MedChemExpress), Gamma-aminobutyric acid (40 mM stock in water, Cat#A2129, Sigma-Aldrich), the GABA receptor antagonist picrotoxin (50 mM stock in DMSO, Cat#P1675, Sigma-Aldrich), Dexamethasone (40 mM stock in DMSO, Cat#D4902, Sigma-Aldrich), and 5-fluorouracil (40 mM stock in DMSO, Cat#F6627, Sigma-Aldrich). Whenever possible, stock solutions were prepared with the mentioned diluent, aliquoted and stored at -80 degrees Celsius.

### NCI diversity set VII and library preparation

The National Cancer Institute Developmental Therapeutics Program (NCI/DTP) https://dtp.cancer.gov provided the NCI Diversity set VII and the extra NSC compounds. The NCI diversity set VII contained 1,581 compounds at 10 mM in DMSO. Compounds were first diluted at 200 µM in water and 20 µL was added into the wells containing 4-day old cell monolayer (final concentration of 20 µM) one-hour prior infection.

### Infection of HT-29 cells and plate fixation

Cell-to-cell spread was determined by infecting a 4-day old confluent monolayer of HT-29 using exponential phase CFPexpressing *S. flexneri* ^20,21^. Briefly, bacteria were grown for 3 hours at 37°C under agitation (180 rpm) from overnight culture at 30°C. Bacteria were diluted in cell media and were added directly to the cell medium. Plates were centrifuged at 1,000 rpm for 5 min at room temperature to initiate and synchronize infection. After one hour of incubation at 37°C in 5% CO_2_ to allow for bacterial invasion, IPTG (10 mM final concentration) and gentamicin (20 µg/ml final concentration) were added to wells to induce the expression of CFP (*S. flexneri*) and to kill extracellular bacteria, respectively. The plate was incubated for 7 hours at 37°C in 5% CO_2_. Infection was stopped after fixation using 4% paraformaldehyde (Electron Microscopy Sciences, cat# 15710) in PBS. The integrity of the cell monolayer was verified using Hoechst staining.

Chemical compounds were added 1-hour prior to infection. A variation is mentioned in the figure legends for conditions where compounds were added 1-hour post-infection, at the same time as IPTG and Gentamicin. Mock condition refers to the DMSO diluent/vehicle concentration used to dissolve compounds.

### Image acquisition and analysis

After fixation, plates were imaged using an ImageXpress Micro imaging system (Molecular Devices). Image analyses were conducted on at least 50 bacterial infection foci in each condition of each biological replicate, if possible. The image analysis was performed with the ImageXpress imaging software (Molecular Devices), as previously described ^82^. Briefly, CFP channel images corresponding to *S. flexneri* were subjected to intensity thresholding (lower threshold set to 1,250 arbitrary units) to segment bacterial signal. Segmented pixels were grouped into objects based on spatial proximity (15pixel connectivity). Objects were filtered to exclude those smaller than 250 au or larger than 8,500 au (usually corresponding to 2 or more foci), as well as those in contact with the image borders (Supplementary Figure 1). The total area of the remaining objects was quantified for each field and exported for downstream analysis. 6 fields per well were acquired and analyzed. The average and number of foci per field are reported in Supplementary. Table 1 and 2. Assay robustness was assessed for each of the twenty screening plates using the Z′-factor ^83^, calculated from the mean and standard deviation of SAR405 (positive control) and mock (DMSO, negative control) wells present on every plate (Supplementary Table 1). Z′ values ranged from 0.066 to 0.713, with 15 plates on 20 falling within the range considered excellent for a cell-based phenotypic assay (Z′ > 0.5) and the remainder within the acceptable range (0 < Z′ < 0.5), indicating robust separation between control conditions on most plates. Because five plates fell below 0.5, all candidate hits, including those identified on plates with comparatively lower Z′ values, were subjected to the same triplicate dose-response confirmation, mitigating the risk of plate-specific false positives. The plate of origin and Z′ value for each primary hit are reported in Supplementary Table 1.

### Hit selection and validation from the phenotypic screen

Screening was performed across twenty 96 well plates containing the 1,581 compounds of the NCI Diversity Set VII, as mentioned in Supplementary Table 1. Each plate contained internal control wells treated with the PIK3C3/hVPS34 inhibitor SAR405, which has been shown to reduce *S. flexneri* dissemination, as well as multiple vehicle-treated control wells ^17^. Following infection and image acquisition as described before, six fields were acquired per well and analyzed using automated image analysis to determine the average infection focus size and the number of infection foci per image. For each well, the mean infection focus size was calculated from the six acquired images. For each individual 96-well plate, the mean and standard deviation of infection focus size values from all compounds were calculated. Hit selection thresholds corresponding to the plate mean minus two standard deviations (mean − 2SD) and mean minus three standard deviations (mean − 3SD) were determined independently for each plate. 61 hits presented an average of infection focus size below these thresholds (Supplementary Table 1). Those hits were subjected to manual curation to eliminate compounds displaying intrinsic autofluorescence, visible precipitation, host cytotoxicity characterized by loss of the epithelial monolayer, or complete absence of infection foci preventing reliable dissemination analysis (Figure 1; Supplementary Figures 1, 2). Following this filtering step, 34 compounds were retained for secondary validation as presented in Supplementary tables 1 and 2.

Fresh powder stocks of the 34 selected compounds were provided by the National Cancer Institute Developmental Therapeutics Program (NCI/DTP) https://dtp.cancer.gov. After resuspension in DMSO, they were retested in secondary screening assays at final concentrations of 5, 10, and 20 µM. Compounds were tested in triplicate wells, with six compounds analyzed per 96-well plate, and the experiment was repeated three time independently. Image acquisition and quantitative analysis were performed as described above and are shown in Supplementary Table 2. Based on reproducibility and dose-dependent reduction in infection focus size, 19 compounds were selected for further characterization (Table 1; Supplementary Table 2).

### Effect on bacterial growth

For growth experiments, overnight cultures in appropriate medium were diluted to an initial optical density at 600 nm of 0.025 in fresh media. Cultures were incubated at 37°C for 3 h (LB) or 5 h (minimal medium) in the presence or absence of compounds as mentioned on figures for end-point measures. For growth curve kinetics, culture tubes (4 mL) were incubated at 37°C, and 300 µL was collected at the indicated time points for measurement. Bacterial growth was quantified by measuring Optical density at 600nm via spectrophotometry (Genesys 20, ThermoScientific). All assays were performed at least three independent times.

During metabolites rescue experiments, medium was supplemented with the corresponding concentration indicated on the graph with adenine (50 mM stock in DMSO, Cat# A2786, Sigma), Guanosine (50 mM stock in water, Cat#G6264, Sigma), Hypoxanthine (50 mM stock in 0.1 M NaOH, Cat#H9636, Sigma), Uracil (50mM stock in 0.1 M NaOH, Cat#U1128, Sigma), Uridine (50 mM stock in water, Cat#U3003, Sigma), Cytidine (50 mM stock in water, Cat#C122106, Sigma), thymidine (50 mM stock in water, Cat#T1895, Sigma) or the corresponding solvent control.

### Cytotoxicity/cell viability

Cell viability was assessed using the CellTiter-Glo Luminescent Cell Viability Assay (Promega), a homogeneous method that quantifies intracellular ATP as an indicator of metabolically active cells. HT-29 cells were seeded in 96-well plates and cultured to form a confluent monolayer over 4 days. Compounds were added at following a semi-log range of 1 to 100 µM, followed by incubation for 1 hour. Subsequently, fresh medium was added to mimic the infection protocol (Gentamicin and IPTG step), and cells were further incubated for either 8 or 24 hours.

For the viability assay, the CellTiter-Glo reagent was prepared according to the manufacturer’s instructions by reconstituting the substrate with the provided buffer and equilibrating to room temperature. Following treatment, culture medium was removed from each well and replaced with serum-free McCoy’s 5A medium (Gibco). An equal volume of CellTiter-Glo reagent was then added to each well. Plates were mixed on an orbital shaker for 2 minutes to induce cell lysis and incubated at room temperature in the dark for 10 minutes to stabilize the luminescent signal. Luminescence was recorded using a plate reader (BioTek Synergy 2).

### Cell death detection (TUNEL assay)

The effect of the selected hits on HT-29 cell death was assayed using the In-situ cell death detection kit, TMR red (Cat#12156792910, Roche), detecting DNA breakage characteristic of certain forms of cell deaths. To reflect the screen, compounds were added at a final concentration of 20 µM on the top of a 4-day HT-29 layer and incubated for 24 h before fixation with PBS+PFA 4% for 15 minutes. The 24 h incubation was used to highlight a long-term effect of these compounds that cannot be detected in a shorter time. The staining was performed according to manufacturer’s recommendations. Three independent biological replicates were performed.

### HT-29 proliferation assay

To test the effect of compounds on the proliferation of actively dividing HT-29 cells, as opposed to the 4-day old monolayer used in infection and viability assays, HT-29 cells were seeded in 96-well plates at 3×10^4^ cells per well and incubated overnight to allow adherence. The day after, the medium is changed with fresh McCoy’s medium containing 10 % FBS and the appropriate compounds were added into the wells and incubated for 15 min (t0.25), 24h, 48h and 72h. After corresponding incubation time, the content of the wells was fixed with DPBS containing 4 % Paraformaldehyde (PFA) for 15 minutes, washed with DPBS three times, stained with 1 µg/mL of Hoechst, in DPBS containing 0.1 % Triton X-100 to permeabilize the cells for ten minutes. Plates were washed three times and maintained in DPBS until imaging. Images were acquired using ImageXpress Micro imaging system using a 10X objective, with six fields acquired per well. Nuclei were counted using the ImageXpress count nuclei module. The average number of nuclei per field was used to generate the growth curve. Experiments were repeated independently at least three times.

### Intracellular CFU recovery

For testing the effect of compounds on intracellular growth, a 4-day cell monolayer was infected with around 2×10^6^ CFU of mid-log *S. flexneri* (MOI around 20). Briefly, after adding medium containing *S. flexneri*, the plate was centrifuged at 1,000 rpm for 5 min to synchronize infection and incubated at 37°C with 5% CO2 for one hour. Gentamicin at 50 µg/mL is added in the medium for one hour to kill extracellular *S. flexneri*. At two hours post-infection, the medium is removed, washed four times with sterile DPBS and fresh cell medium is added to the well with compounds and a lower dose of gentamicin (20 µg/mL) for four extra hours (6 hpi) or content of at least 6 wells were lysed to verify. At the appropriate time point, infected cells were washed three times with DPBS to eliminate gentamicin and compounds, lysed with 0.5 % Triton X-100 in DPBS, diluted and plated on LB agar plates containing Congo Red. After overnight incubation at 37°C, red CFU on plates were counted to estimate the number of CFU recovered from well. The number of CFU corresponds to the average of three wells. The experiment was repeated independently three more times.

### Statistical analysis

Statistical analyses of the values were conducted using the GraphPad Prism 9.0.2 software. Analysis of a single variable across two groups was performed with unpaired t-test to determine statistical significance. Experiments with multiple groups were analyzed with one-way ANOVA followed by Dunnett’s multiple comparisons. P-values were represented on figures as follows: ns, not significant; *, p<0.05; **, p<0.01; ***, p<0.001; ****, p<0.0001. All values used for statistical analysis represent the average of at least three independent experiments.

### Data availability

All data supporting the findings of this study are available within the paper and its Supplementary Information, and from the corresponding author upon reasonable request. Figures and panels made using biorender.com are accessible using link mentioned on the legend.

## Acknowledgments

We wish to thank the National Cancer Institute Developmental Therapeutics Program (NCI/DTP) https://dtp.cancer.gov for providing compounds present in this study. We thank all the members of the Agaisse laboratory for discussions on the project. This study was funded by NIH NIAID R01 AI153455 (H. Agaisse, D. Engel) and R01AI179778 (H. Agaisse). The funder played no role in study design, data collection, analysis and interpretation of data, or the writing of this manuscript.

## Author contributions

S.R, T.M, D.E and H.A conceived and designed the experiments. S.R, T.M and E.D performed experiments and acquired data. S.R and H.A analyzed and interpreted the data. S.R generated the figures. S.R and H.A wrote the first draft of the manuscript. H.A and D.E acquired funding and supervised the project. All authors reviewed and edited the manuscript.

## Competing interests

All authors declare no financial or non-financial competing interests.

## Funding

This work was supported by the National Institutes of Health grants R01AI073904 (H.A.), R01AI153455 (H.A. and D.E.) R03AI142654 (D.E.).

## Supplementary Figures

**Supplementary Fig 1:**
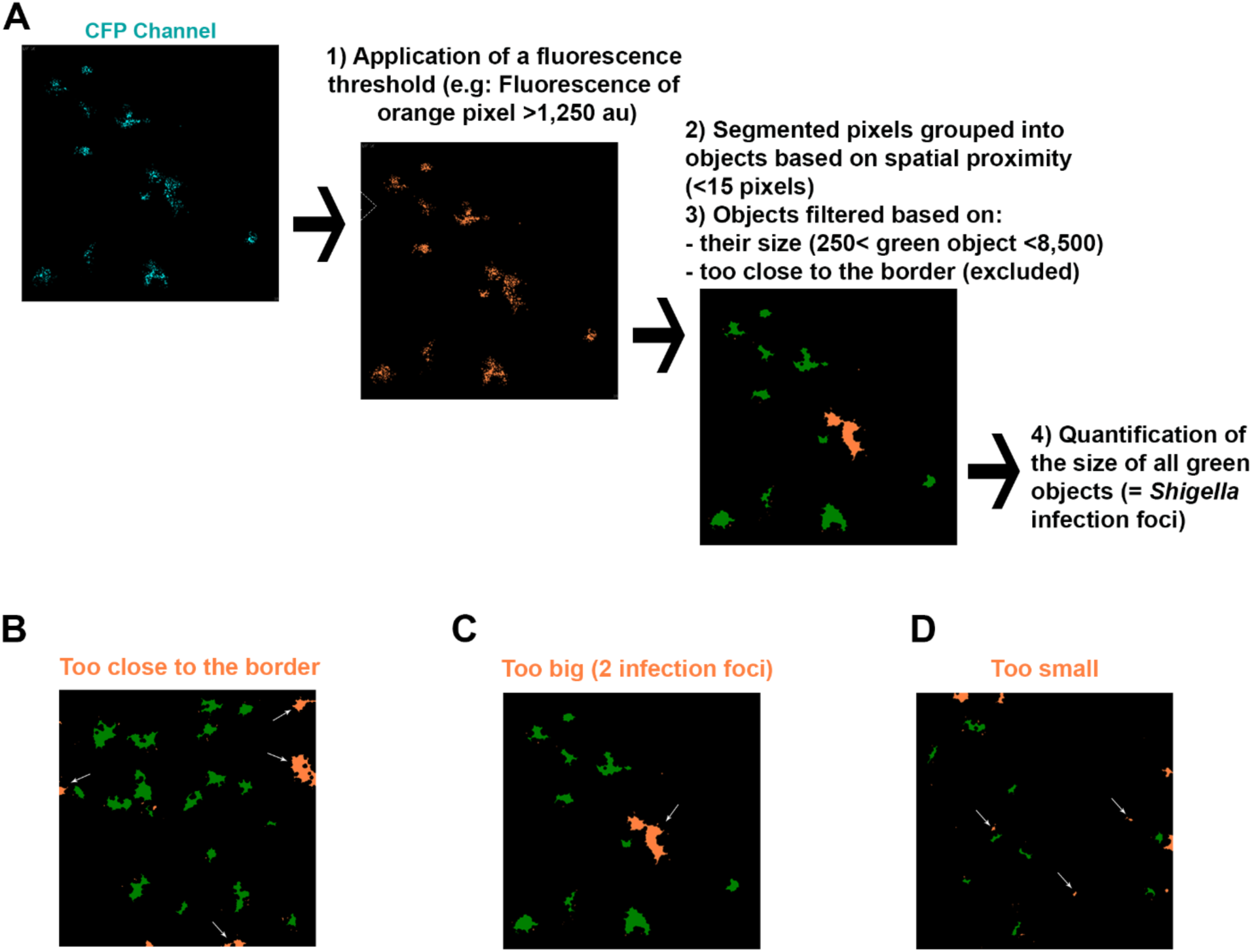
MetaXpress software analysis. **(A)** Steps performed by the software MetaXpress to create and quantify object size based on CFP channel image (Channel image (*Shigella*). **(B-D)** Exemple of exclusion based on the location of object on the border of the field (B), and the size if too big (C) or too small (D).

**Supplementary Fig 2:**
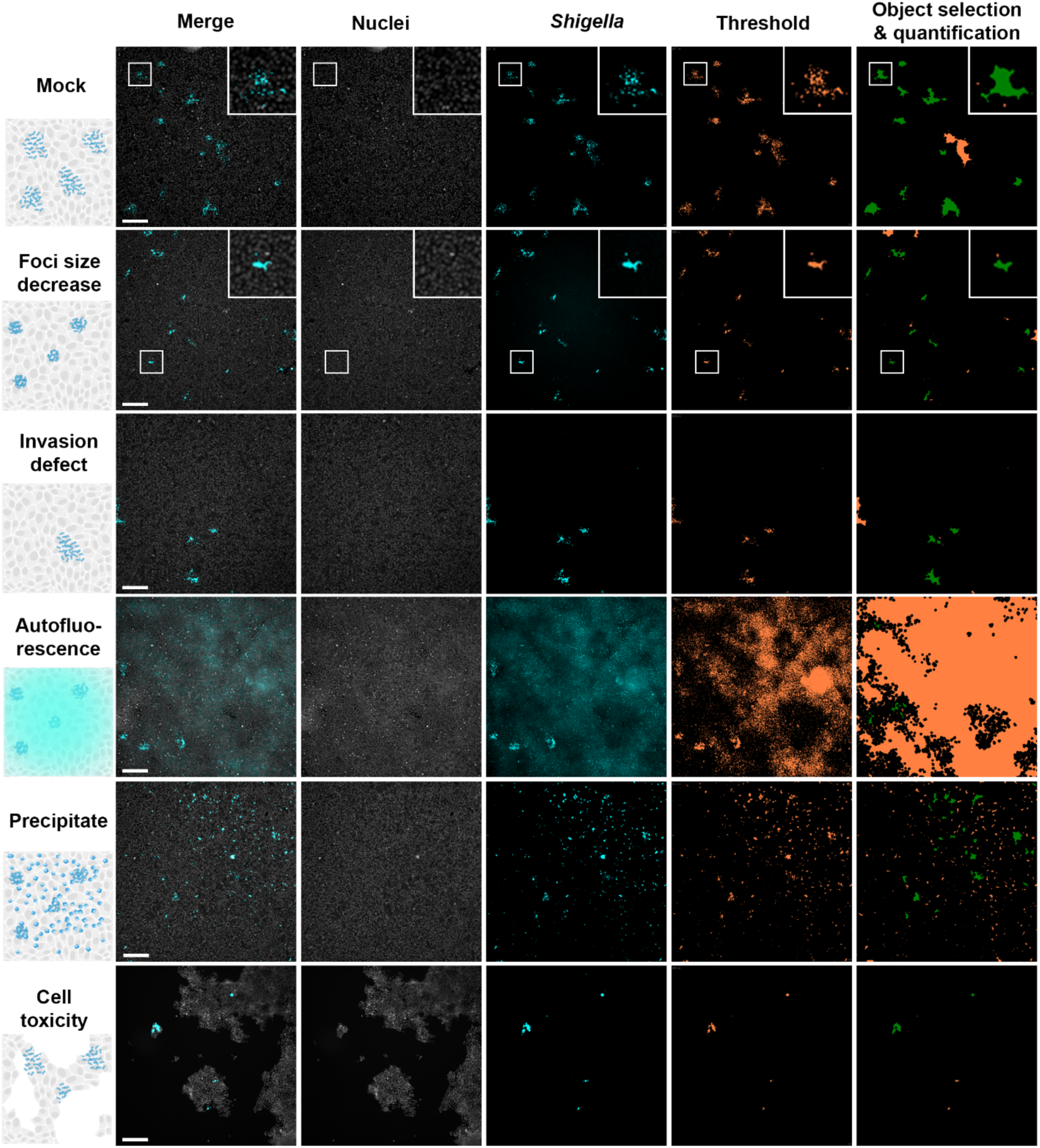
Representative images of each phenotype observed and mentioned in Figure 1. The presence of precipitation or autofluorescence does not permit an accurate quantification of the Infection foci. A compound with strong cytotoxicity, as shown by the lack of a full monolayer, is excluded from the analysis. Nuclei were stained using Hoescht to visually control the state of the monolayer. Scale bar is 100 µm.

**Supplementary Fig 3:**
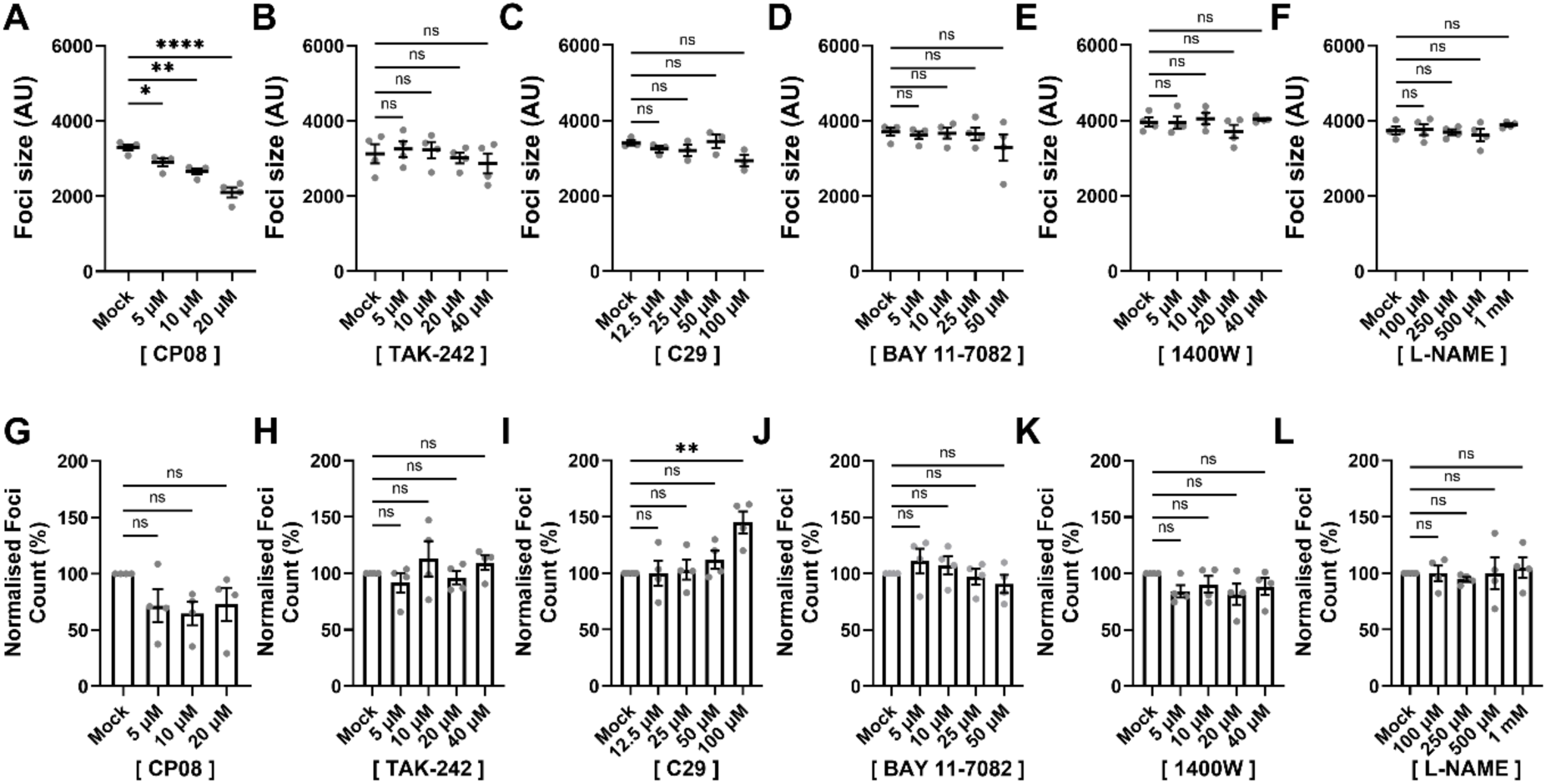
CP08 (NSC126224) restricts *Shigella* dissemination independently of canonical TLR signaling. **(A-F)** Quantification of *Shigella* infection foci size in arbitrary units 8 hours post-infection in the presence of different concentrations of CP08 (A), the TLR4 inhibitor TAK-242 (B), the TLR2 inhibitor C29 (C), the NF-κB pathway inhibitor BAY 11-7082 (D), inducible nitric oxide synthase (iNOS) inhibitor 1400W (E), and the nitric oxide synthase (NOS) inhibitor L-NAME (F). **(G-L)** Quantification of *Shigella* infection foci number normalized on Mock condition in the presence of different concentrations of CP08 (G), the TLR4 inhibitor TAK-242 (H), the TLR2 inhibitor C29 (I), the NF-κB pathway inhibitor BAY 11-7082 (J), inducible nitric oxide synthase (iNOS) inhibitor 1400W (K), and the nitric oxide synthase (NOS) inhibitor L-NAME (L). Each dot represents one biological replicate. Error bars, standard error of the mean; statistical analysis, one-way ANOVA with Dunnett’s multiple comparisons test; ns, not significant; *, p<0.05; **, p<0.01; ****, p<0.0001.

**Supplementary Fig 4:**
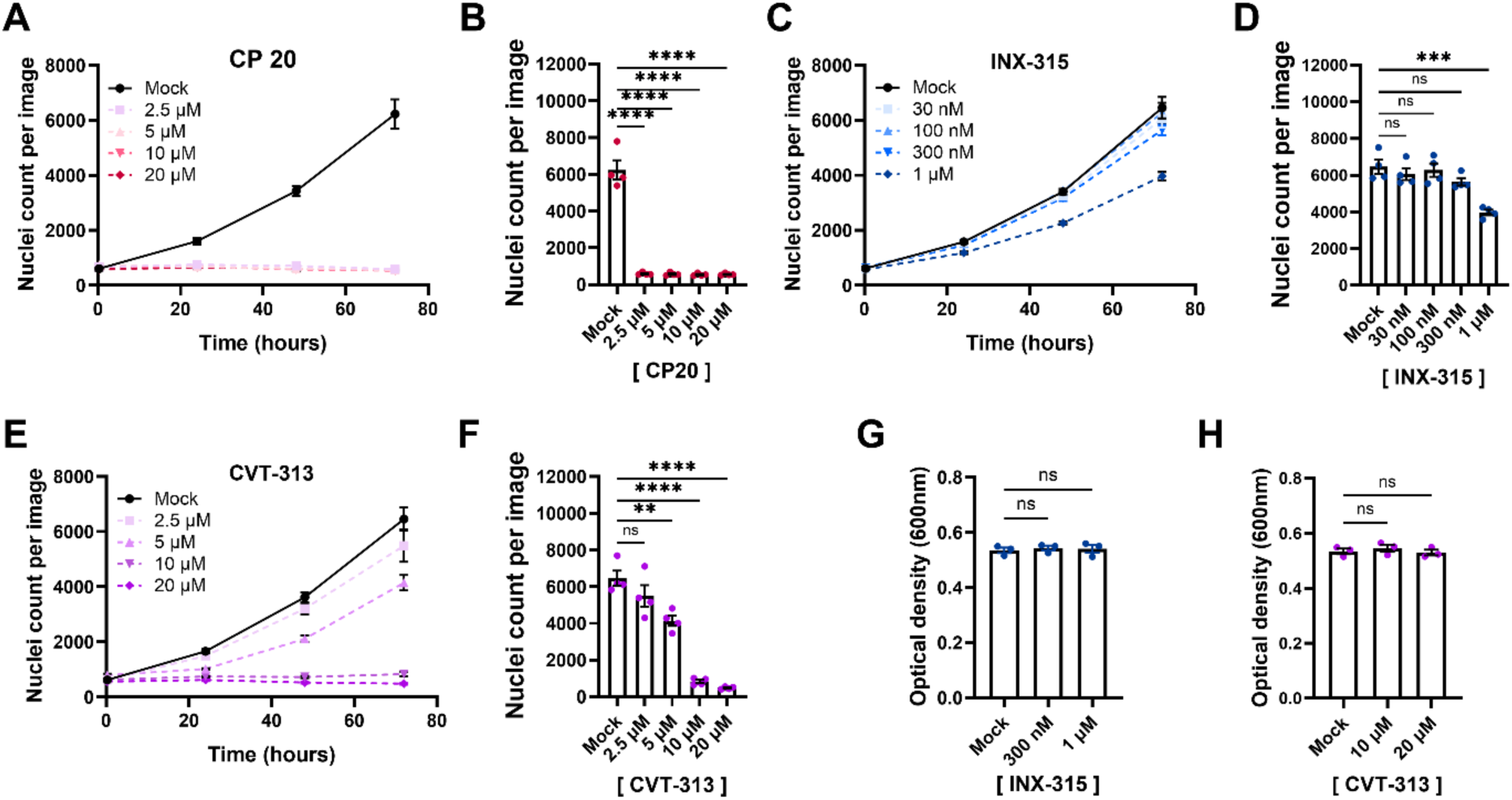
CP20, INX-315 and CVT-313 inhibit HT-29 cell growth. **(A-B)** Effect of different concentrations of CP20 on HT-29 cell growth represented as curves (A) or the count of nuclei per image at 72h (B), in presence of different concentrations of CP20. (**C-D)** Effect of different concentrations of INX-315 on HT-29 cell growth represented as curves (C) or the count of nuclei per image at 72h (D). **(E-F)** Effect of different concentrations of CVT-313 on HT-29 cell growth represented as curves (E) or the count of nuclei per image at 72h (F). **(G-H)** Optical density at 600nm after 3h growth in LB of *Shigella flexneri* in presence of INX-315 (G) and CVT-313 (H). Each dot represents one biological replicate. Error bars, standard error of the mean; statistical analysis, one-way ANOVA with Dunnett’s multiple comparisons test; ns, not significant; **, p<0.01; ***, p<0.001; ****, p<0.0001.

**Supplementary Fig 5:**
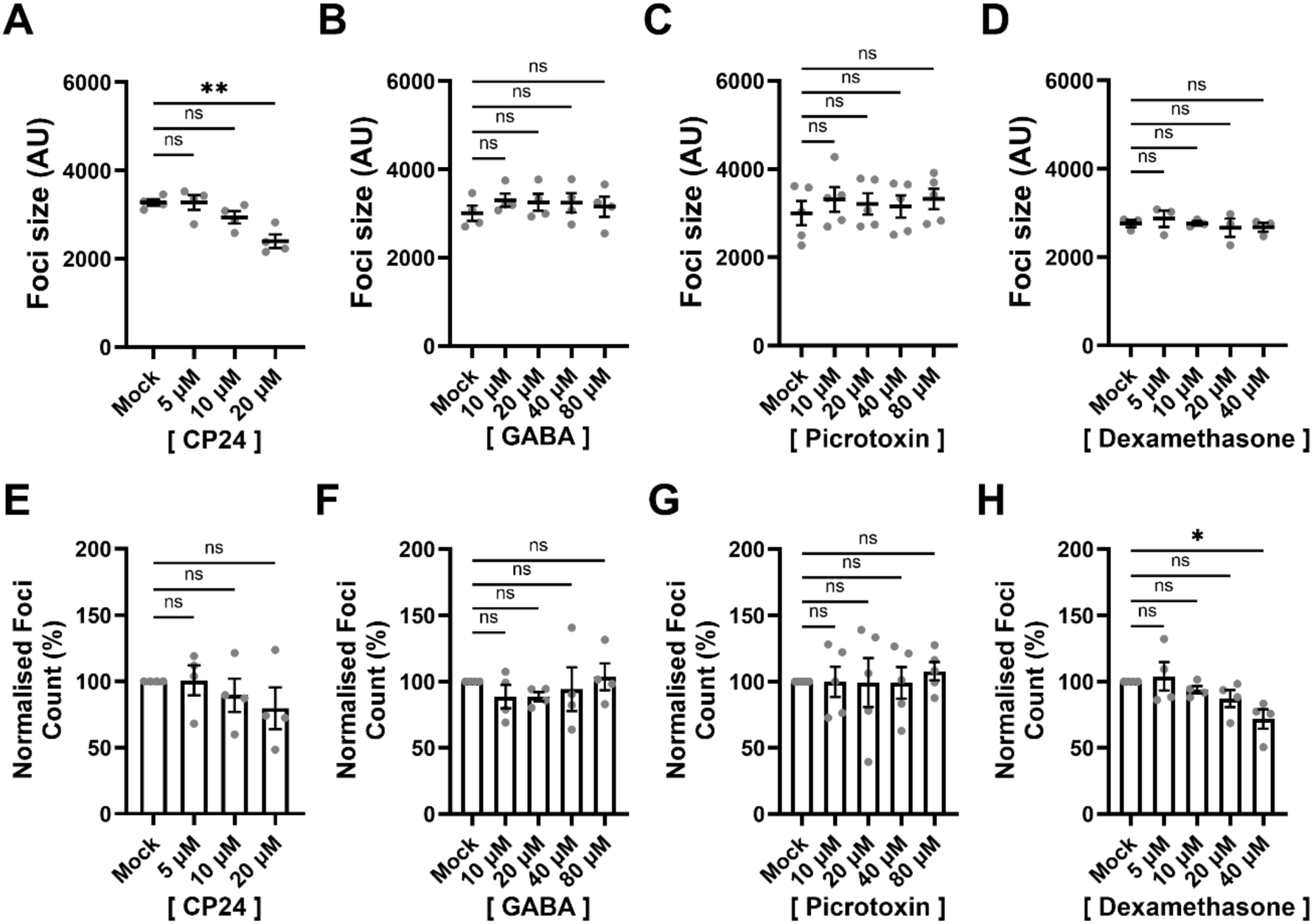
CP24 (NSC70799) reduces *Shigella* infection foci independently of neurosteroid and glucocorticoid signaling. **(A-D)** Quantification of *Shigella* infection foci size in arbitrary units 8 hours post-infection in the presence of different concentrations of CP24 (A), the GABA(A) receptor agonist Gamma-aminobutyric acid (B), the GABA(A) receptor antagonist Picrotoxin (C), and the glucocorticoid receptor agonist Dexamethasone (D). **(E-H)** Quantification of *Shigella* infection foci number normalized on Mock condition in the presence of different concentrations of CP24 (E), the GABA(A) receptor agonist Gamma-aminobutyric acid (F), the GABA(A) receptor antagonist Picrotoxin (G), and the glucocorticoid receptor agonist Dexamethasone (H). Each dot represents one biological replicate. Error bars, standard error of the mean; statistical analysis, one-way ANOVA with Dunnett’s multiple comparisons test; ns, not significant; *, p<0.05; **, p<0.01; ****, p<0.0001.

## Notes

### Competing Interest Statement

The authors have declared no competing interest.

